# Compact serum miRNA qPCR model for pancreatic cancer discrimination with independent and clinical validation

**DOI:** 10.64898/2026.05.11.724428

**Authors:** Sunao Yotsutsuji, Hiroki Kataoka, Tomomi Ando, Mika Inada, Mitsuko Sugano, Masaaki Takada, Minoru Esaki, Ken Kato, Yusuke Yamamoto, Yoshitake Sano

**Author notes:** These authors contributed equally (Sunao Yotsutsuji, Yoshitake Sano).

## Abstract

**Background:** For pancreatic cancer, practical blood-based tests for early detection and postoperative surveillance remain elusive. We sought to develop a qPCR-measurable serum microRNA (miRNA) panel that robustly discriminates pancreatic cancer from non-cancer controls and other malignancies.

**Methods:** We profiled 255 serum miRNAs in batch 1 (n=72) and selected 27 candidates. Candidates were refined in batch 2 (n=552) and cross-batch evaluation was performed with batch 3 (n=391) to derive a miRNA model. Independent validation used batch 4 (n=616). Clinical relevance was assessed in an independent clinical cohort of resection patients with samples obtained preoperatively and at 1 and 12 months postoperatively.

**Results:** The miRNA model trained on batches 2 and 3 achieved an area under the curve (AUC) of 0.91 and 0.83 for pancreatic cancer versus non-cancer controls and non-cancer plus other cancers, respectively, when independently validated in batch 4. Stage-wise AUCs in batch 4 were 0.91 (I), 0.94 (II), 0.86 (III) and 0.90 (IV). In the clinical batch, the score decreased postoperatively (preoperative vs month 1; p<0.01) and was higher in recurrence than non-recurrence (p<0.001).

**Conclusions:** The developed compact miRNA qPCR assay discriminated pancreatic cancer across independent assay batches and showed clinical relevance for postoperative surveillance.

**Clinical Trial Registration:** Not applicable.

## Background

Pancreatic cancer (PC) remains one of the most lethal malignancies, with a 5-year survival rate below 10%, largely because most patients are diagnosed at an advanced stage^1^. Early diagnosis is critical for improving outcomes; however, PC is frequently asymptomatic in its early stages, and when do symptoms occur, they are often nonspecific^2^. In addition, the pancreas is located deep within the abdominal cavity, which limits the feasibility of physical examination and broad imaging-based screening^3^. Conventional blood-based biomarkers such as carbohydrate antigen 19-9 (CA19-9) have limited sensitivity and specificity, particularly for early-stage disease and for differentiating PC from benign pancreatic disorders such as chronic pancreatitis^4–6^. These challenges highlight the need for practical blood-based approaches that can support detection at a potentially curable stage and aid clinical management. Moreover, there are few effective blood-based tools for postoperative surveillance and early identification of recurrence after curative-intent resection.

Circulating microRNAs (miRNAs) are promising minimally invasive biomarkers for cancer detection and monitoring^7,8^. miRNAs are small non-coding RNAs that regulate gene expression at the post-transcriptional level, and their dysregulation is linked to oncogenic processes^9,10^. Importantly, miRNA profiles can change early during tumour development, potentially reflect tumour-related molecular alterations^11^. Circulating miRNAs are also relatively stable in serum and plasma because they are protected by extracellular vesicles and RNA-binding proteins, making them amenable to blood-based testing^12^. Taken together, these features make miRNAs attractive candidates for developing diagnostic assays that are scalable and suitable for routine measurement.

High-throughput platforms such as next-generation sequencing and microarrays have enabled comprehensive discovery of PC–associated miRNAs, revealing signatures that discriminate cancerous from non-cancerous samples^11,13^. Several studies have combined omics-scale measurements with machine-learning models and reported high discrimination performance^13–15^. Nevertheless, translation into routine clinical testing remains limited. High-throughput assays often require careful consideration of cost, technical complexity, and bioinformatics resources. Furthermore, variability arising from technical factors such as batch effects has been reported for some published signatures, underscoring the need for rigorous validation to support reproducibility and generalisability across independent assay batches and laboratories^8,16^. From a clinical perspective, robust discrimination of PC from other common malignancies is also essential, yet comparatively few studies have systematically evaluated performance in realistic settings that include multiple cancer types.

Quantitative PCR (qPCR) is widely available in clinical and research laboratories and provides a practical platform for implementing miRNA-based assays. However, reducing candidate biomarkers to a minimal panel can increase vulnerability to technical noise and batch-specific effects unless robustness is explicitly addressed. In addition to conventional normalisation, feature engineering strategies that mitigate inter-batch variability can help to improve model stability without requiring complex workflows. Therefore, a compact, analytically stable miRNA panel that maintains performance across independent assay batches and against other malignancies would represent an important step toward clinically deployable PC testing.

In this study, we developed a qPCR-measurable serum miRNA model to discriminate PC from non-cancer control (NC) and other malignancies, using a multi-phase design with cross-batch evaluation and independent validation. A compact miRNA model showed high discrimination of PC from NC and retained performance when other malignancies were included. We further assessed the clinical relevance for postoperative surveillance in an independent National Cancer Center biobank cohort, where the score decreased after surgery and differed between the recurrence and non-recurrence groups during postoperative follow-up.

## Material and Methods

### Serum samples and ethics

All experimental procedures for the analysis of commercial serum samples were reviewed and approved by the Gene Analysis Research Committee of the Corporate Research & Development Center, Toshiba Corporation. Experiments were conducted in compliance with the institution’s Regulations for Human Research. Serum samples for model development and validation (independent assay batches 1–4) were purchased from ProteoGenex (Culver City, CA), a commercial biobank operating under institutional review board-approved protocols, and stored at −80 °C until miRNA extraction.

For the clinical cohort, serums from each subject were obtained from the National Cancer Center (NCC) biobank, and the clinical data was corrected by electrical health record under an institutional review board-approved protocol entitled “Development of recurrence markers based on serum miRNA concentrations before and after surgery for pancreatic cancer” (National Cancer Center Research Ethics Review Committee; approval number: 2024-023). Written informed consent had been obtained from all participants for the use of biospecimens in research. The study was conducted in accordance with the Declaration of Helsinki. Only the subset of specimens and analyses corresponding to the results presented in this manuscript were included.

### Study design and independent assay batches (marker selection)

This retrospective study aimed to develop a serum-based miRNA panel for detection of PC. The workflow comprised four sequential phases: (1) discovery and initial marker selection (batch 1), (2) candidate refinement (batch 2), (3) model development with cross-batch validation (batch 3), and (4) independent validation of the final model (batch 4) (Fig. 1).

Independent assay batches were constructed as follows.
Batch 1 (discovery; Group 1): A total of 72 serum samples from individuals with PC, breast cancer, and NC were analysed by profiling 255 miRNAs.
Batch 2 (candidate refinement; Group 2): A total of 552 samples, including pancreatic, breast, lung, gastric, and colorectal cancers as well as NC, were analysed for the 27 candidate miRNAs, and 13 miRNAs contributing to discrimination of PC were identified.
Batch 3 (model development and cross-batch validation; Group 3): A total of 391 samples comprising the same cancer types as batch 2 were analysed for the 13 miRNAs. A discrimination model demonstrating consistent performance across batches 2 and 3 was developed and further refined to a compact miRNA panel.
Batch 4 (independent validation): A total of 616 newly collected serum samples were analysed for the miRNAs, and the final model was independently validated.
A summary of the sample numbers by cancer type, disease stage, sex, and age is provided in Supplementary Table 1.

**Fig. 1:**
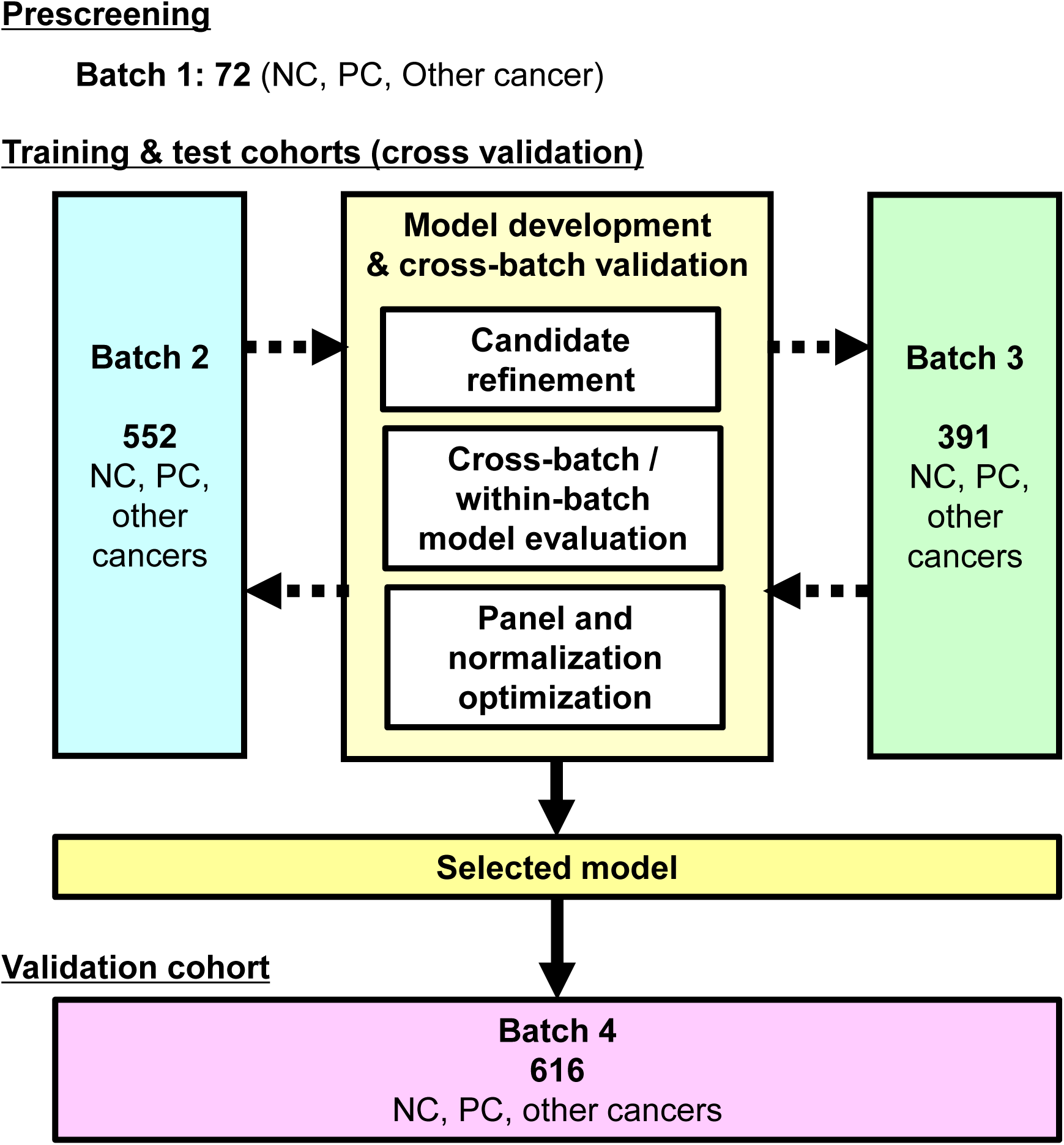
Study design for serum miRNA model development and independent validation. Schematic overview of the multi-phase model discovery, development, and validation workflow. Batch 1 (n = 72) was used for discovery profiling and initial candidate identification. Batches 2 (n = 552) and 3 (n = 391), which included non-cancer control (NC), pancreatic cancer (PC), and other cancer types, were used for cross-batch and within-batch evaluation. The final model was independently validated using Batch 4 (n = 616), which included NC, PC, and other cancer types.

### Clinical cohort and endpoints

The eligibility criteria for the NCC cohort were as follows: (i) patients with PC who underwent surgery at the National Cancer Center (NCC) between 1 January 2018 and 30 April 2024; (ii) age ≥18 years; (iii) broad consent for biobank research use; (iv) R0 resection; and (v) availability of archived serum at prespecified perioperative and follow-up time points. The recurrence status was determined based on clinical assessment supported by imaging and/or pathological findings over a 1-year follow-up. For the analyses, we evaluated (i) the perioperative change in the miRNA-derived score between the preoperative and postoperative samples obtained at 1 month postoperatively, and (ii) the recurrence status in samples obtained at 12 months (Supplementary Table 2).

### miRNA extraction

miRNA was extracted from 300 µL of serum using the NucleoSpin miRNA kit (Macherey-Nagel, Düren, Germany) according to the manufacturer’s protocol. RNA was eluted in 30 µL of RNA Storage Solution (Thermo Fisher Scientific, Waltham, MA) and stored at −80 °C until use.

### Quantitative PCR

Complementary DNA (cDNA) was synthesised from 2 µL of miRNA extract using the TaqMan™ Advanced miRNA cDNA Synthesis Kit (Thermo Fisher Scientific; Cat. A28007) following the manufacturer’s instructions. qPCR was performed using TaqMan™ Advanced miRNA Assays (Thermo Fisher Scientific; Cat. A25576) and TaqMan™ Fast Advanced Master Mix (Thermo Fisher Scientific; Cat. 4444964) according to the manufacturer’s protocols. The IDs of the TaqMan probes used and the corresponding target miRNA sequences are listed in Supplementary Table 3. All reactions were conducted in technical triplicates. Quantitative values were derived from the threshold cycle (Ct) number.

In batch 1, relative expression levels were calculated using the ΔCt method with miR-486-5p as the reference. In batches 2–4 and clinical study, absolute quantification was performed using a standard curve generated from synthetic miRNA oligonucleotides at concentrations of 0, 5 × 10^2^, 1 × 10^3^, 5 × 10^3^, 5 × 10^4^, 5 × 10^5^, and 5 × 10^6^ copies/µL. The synthetic miRNA oligonucleotides were designed based on the miRNA sequences listed in Supplementary Table 2 and were synthesised by 5′-phosphorylation. Values below the lower limit of quantification (500 copies/µL) were assigned as undetected (UD) measurements.

### Construction of the pancreatic cancer discrimination model (screening model)

Batches 1–4 were used for the construction and validation of the screening model. Batch 1 was used for the initial feature selection, batch 2 for the candidate refinement, batches 2 and 3 for the model development and cross-batch validation, and batch 4 for the independent validation of the final model. Expression levels of target miRNAs were normalised to the concentration of miR-486-5p. Two types of features were generated: (1) raw ratios (target miRNA/miR-486-5p) and (2) log10-transformed ratios. All features were standardised as z-scores before model construction. Logistic regression was performed to discriminate PC from NC and other malignancies. The model was implemented using scikit-learn (ver. 1.2.2) with the following hyperparameters: penalty = L2, C = 1,000, and solver = lbfgs.

First, batch 1 was used for the initial feature selection. From the 255 miRNAs profiled in batch 1, miRNAs that showed statistically significant differential expression between PC and NC, or between breast cancer and NC were considered for selection as candidate features. Based on this screening process, 27 miRNAs were selected for subsequent analyses.

Next, batch 2 was used for candidate refinement. The 27 candidate miRNAs were profiled in batch 2 and evaluated for their ability to discriminate PC from NC and other malignancies. Candidate selection consisted of two sequential filtering steps: (i) quality-based filtering and (ii) feature-importance filtering using a gradient boosting decision tree (GBDT). In the quality-based filtering step, miRNAs with a UD ratio >5% or a coefficient of variation (CV) >200% in PC samples were excluded. In the feature-importance filtering step, a GBDT model trained on batch 2 samples was used to classify PC, NC, and other malignancies, and miRNAs with low feature importance were excluded in a similar manner to previous studies^15^. GBDT was implemented using xgboost (ver. 2.1.1) with hyperparameters optimised using Optuna (ver. 4.0.0). Through these sequential filtering steps, 13 miRNAs were selected for downstream analysis.

Next, the models were reconstructed using the selected 13 miRNAs and evaluated using two validation strategies: (a) cross-batch validation, in which models were trained on batch 2 and tested on batch 3 and vice versa; and (b) within-batch validation, based on 10-fold cross-validation using the combined dataset from batches 2 and 3. Cross-batch validation provided a stringent assessment of model generalisability across batches, whereas within-batch validation estimated performance in a pooled-data setting. To identify an optimal compact panel, all possible combinations of five miRNAs from the 13 selected miRNAs were exhaustively evaluated. For each combination, the discrimination models described above were constructed and assessed using both validation strategies by performing receiver operating characteristic (ROC) curve analysis and calculating the area under the curve (AUC) for the discrimination of PC from NC and other malignancies. The final model comprised miR-29c-3p, miR-205-5p, miR-34a-5p, miR-483-5p, miR-324-3p, and miR-486-5p, with expression levels normalised using the geometric mean of these miRNAs. In addition, cel-miR-39-3p was spiked in as an exogenous control in order to monitor the RNA extraction efficiency, and only samples that satisfied the predefined quality criteria based on the spike-in control were included in the analysis.

Finally, the selected model was independently validated using batch 4 samples. Predictions for batch 4 were generated using the model trained on batches 2 and 3, and performance was assessed by calculating the AUC for the discrimination of PC from NC and other malignancies. This independent validation provided an unbiased assessment of model performance in a new sample set and confirmed its potential utility for detecting PC. To assess its potential utility for early-stage detection, model performance against NC was also evaluated stratified by clinical stage in batch 4.

### Prognostic model and score calculation

For the clinical analyses, a separate discrimination model was constructed using the data employed in the screening phase. The model was designed to evaluate whether the miRNA panel selected during screening could distinguish a clinical state with PC from a state without PC. In this context, non-cancer samples used in the screening dataset were regarded as representing the non–pancreatic cancer state. The model was trained using batches 2–4 and was restricted to PC and non-cancer samples. Logistic regression was used as the discrimination algorithm, with regularisation parameters optimised using batches 2–4. The same selected miRNAs used in the screening model were used, together with geometric mean normalisation across these marker values. The fitted model was first evaluated for its ability to discriminate PC from non–pancreatic cancer samples in batches 2–4 and was subsequently applied to the clinical samples to calculate a discrimination score for each sample. This score was used in subsequent analyses to assess perioperative changes and associations with recurrence risk.

### Statistical analysis

Discrimination ability in batches 2–4 was assessed using ROC curves and AUC. For the miRNAs evaluated in batches 2 and 3, performance was assessed using predictions generated by a leave-one-out cross-validation procedure within each batch. For the final model, performance was assessed using predictions generated by a model trained on batches 2 and 3.

For the clinical analyses, perioperative changes (preoperative to postoperative month 1) were evaluated using the Wilcoxon signed-rank test on paired samples. To evaluate postoperative changes in the discrimination score associated with recurrence status, comparisons according to recurrence status were performed at postoperative month 12 samples using the Mann-Whitney U test. For both statistical tests, a p value <0.05 was considered statistically significant unless otherwise specified. Analyses were performed in Python (ver. 3.11).

## Results

### Study overview and batches

We developed and validated a serum miRNA–based discrimination model for PC through a four phase retrospective workflow (Fig. 1). Batch 1 (discovery; n=72) was used for broad profiling and initial marker selection. Batch 2 (candidate refinement; n=552) and batch 3 (model development and cross-batch validation; n=391) included NC, PC, and other malignancies (breast, lung, gastric, and colorectal cancers) and were used to develop and rigorously evaluate the reproducibility and robustness of the candidate models. Batch 4 (independent validation; n=616) consisted of newly collected samples with the same diagnostic categories and was used for independent testing of the final model. Summary characteristics for all batches are provided in Supplementary Table 1.

### Discovery profiling identified 27 candidate miRNAs

In batch 1, we profiled 255 circulating miRNAs using TaqMan PCR and quantified the relative expression levels as ΔCt values normalised to miR-486-5p (Supplementary Table 4). From these profiled miRNAs, candidates showing statistically significant differential expression between PC and NC, or between breast cancer and NC, were considered during the feature selection process. Based on this screening, 27 miRNAs were selected as candidate features for subsequent analyses. Breast cancer samples were included in the discovery batch to facilitate assessment of whether observed miRNA alterations were potentially enriched in PC rather than reflecting general cancer-associated changes.

Next, we quantified the 27 candidate miRNAs in batch 2 by absolute qPCR and normalised each miRNA to miR-486-5p (summary statistics in Supplementary Table 5). Figure 2A shows a heatmap of miRNA expression levels. Unsupervised visualisation using dimensionality reduction did not clearly discriminate among NC, PC, and other malignancies (Fig. 2B). Therefore, we constructed supervised discrimination models and evaluated the discrimination performance for PC. Using the 27-miRNA panel, the model discriminated PC from NC with an AUC of 0.91 (Fig. 2C) and discriminated PC from NC plus other malignancies with an AUC of 0.79 (Fig. 2D). To reduce the panel size while preserving performance, we applied sequential filtering: (i) quality-based filtering excluding miRNAs with an undetected (UD) ratio >5% or high within-PC variability (CV >200%), followed by (ii) feature-importance filtering using an XGBoost model trained in batch 2 to classify PC vs NC vs other malignancies. Through these steps, 13 miRNAs were selected for downstream evaluation (Fig. 2E).

**Fig. 2:**
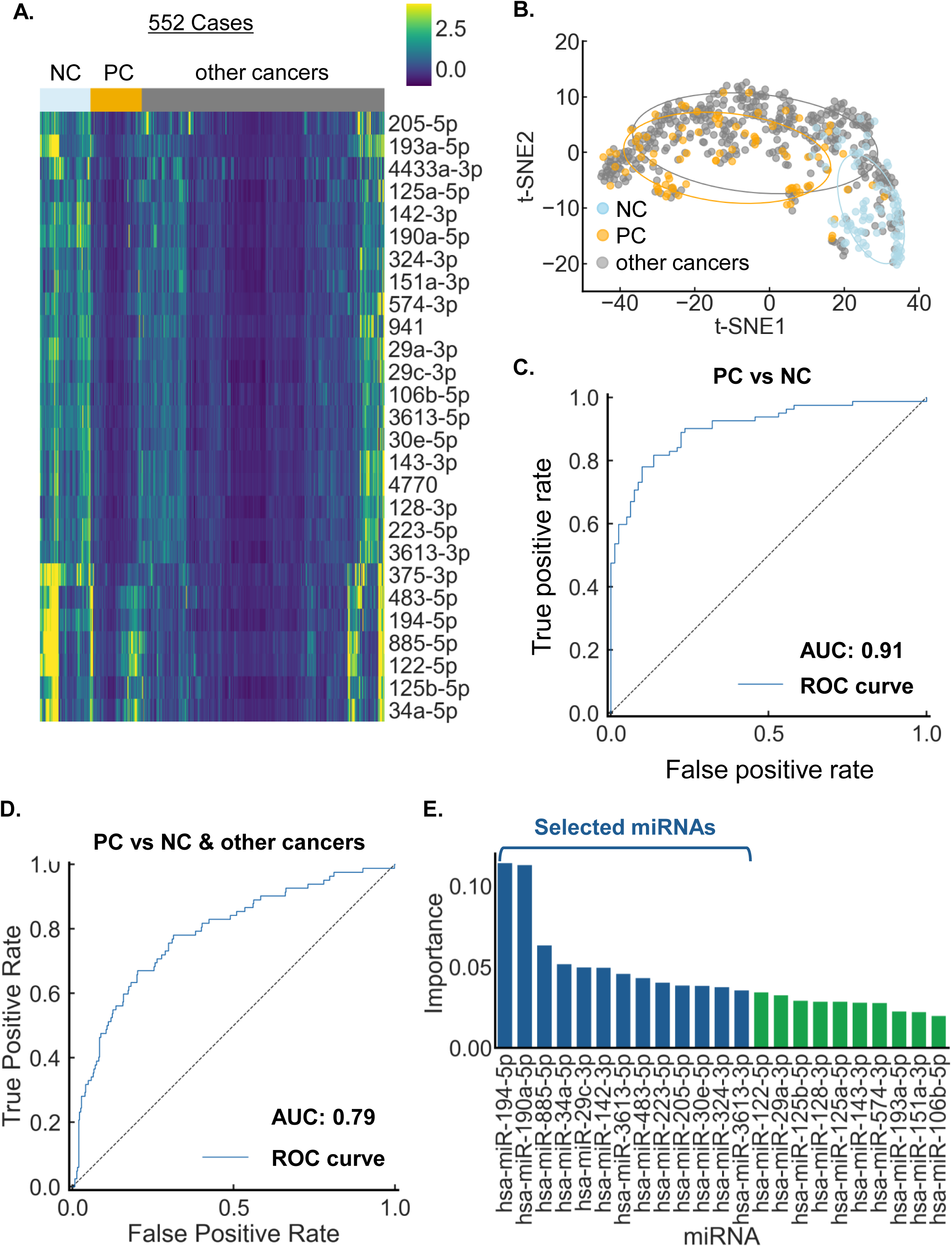
Candidate miRNA profiling in Batch 2 and marker selection. (A) Heatmap of the 27 candidate serum miRNAs quantified in Batch 2 (n = 552; NC, PC, and other cancer types), shown as scaled expression values. (B) Two-dimensional t-SNE embedding of the 27 miRNA expression measurements. (C, D) ROC curves evaluating the discriminatory performance of the 27 miRNAs for (C) PC versus NC and (D) PC versus NC plus other cancer types. (E) Feature-importance ranking of the 23 miRNAs s after initial quality-based filtering, used to select a reduced marker panel. AUC, area under the curve; NC, non-cancer control; PC, pancreatic cancer; ROC, receiver operating characteristic.

### A 13-miRNA panel generalised to batch 3, enabling selection of a compact miRNA panel

We measured the selected 13 miRNAs in batch 3 using the same absolute qPCR and normalisation workflow. The expression patterns are shown in Fig. 3A (summary statistics in Supplementary Table 6). In batch 3, the 13-miRNA panel achieved an AUC of 0.81 for PC versus NC (Fig. 3B) and an AUC of 0.74 for PC versus NC plus other malignancies (Fig. 3C), indicating that the panel retained meaningful discrimination in an independent assay batch, although further improvement appeared necessary for more robust classification.

**Fig. 3:**
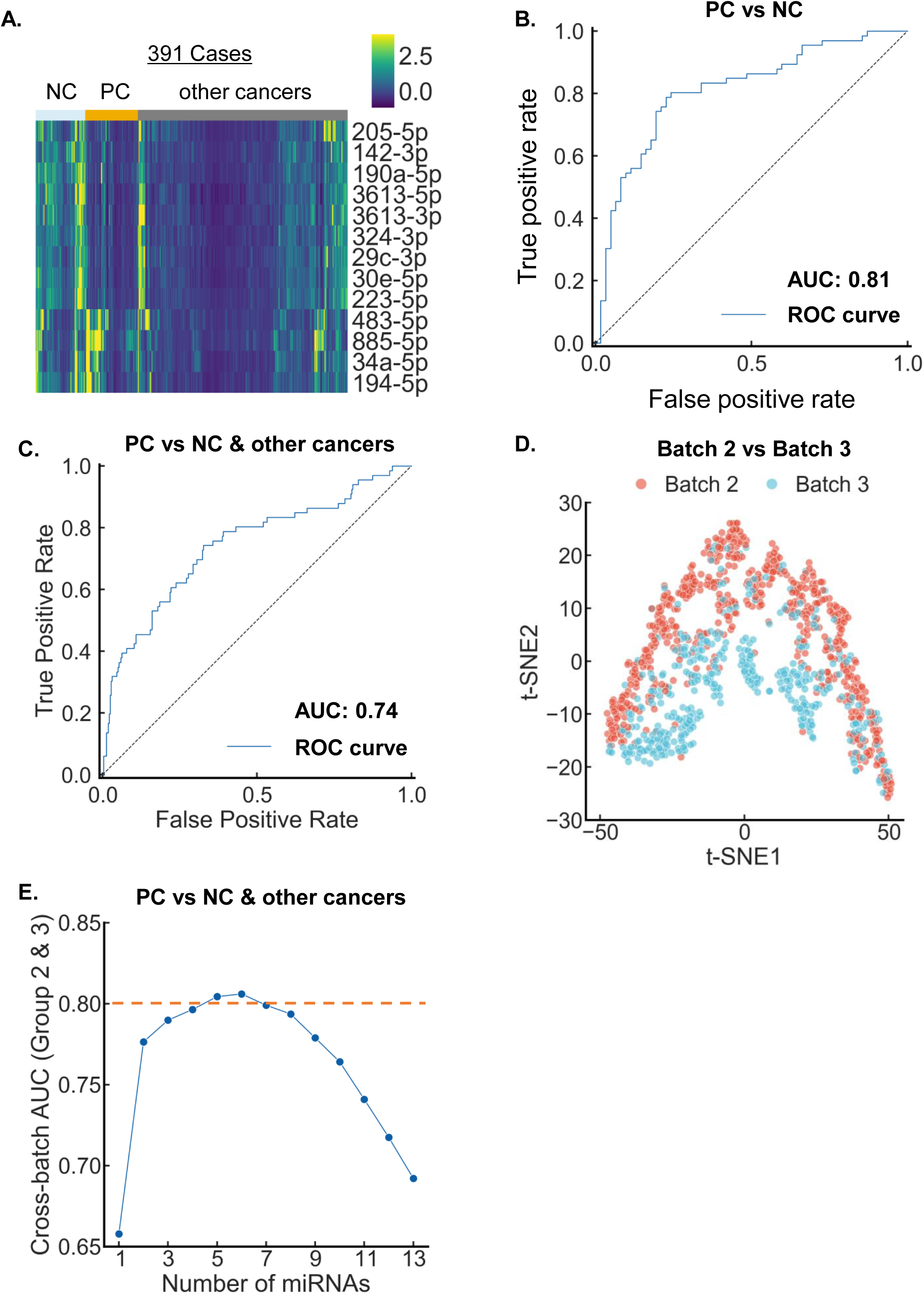
miRNA profiling in Batch 3 and performance of a 13-miRNA panel. (A) Heatmap of the selected 13 serum miRNAs measured in Batch 3 (n = 391; NC, PC and other cancer types) shown as scaled expression values. (B, C) ROC curves in Batch 3 evaluating the performance of the selected miRNA panel for (B) PC versus NC and (C) PC versus NC plus other cancer types. (D) t-SNE visualisation of 13 miRNA expression profiles from Batches 2 and 3. (E) Relationship between panel size and cross-batch discriminatory performance for PC versus NC plus other cancer types. AUC, area under the curve; NCs, non-cancer controls; PC, pancreatic cancer; ROC, receiver operating characteristic.

To explore potential sources limiting cross-batch performance, we assessed batch-to-batch differences between batches 2 and 3. In the unsupervised visualisation of the 13-miRNA measurements using dimensionality reduction, samples tended to separate by assay batch, indicating systematic batch effects (Fig. 3D). Next, we examined how the number of markers affects discrimination performance for PC versus NC plus other malignancies. Using the standard miR-486-5p ratio normalisation, subsets of five to six target miRNAs achieved AUC values above 0.80, whereas the full marker set did not exceed an AUC of 0.80 for this comparison across batches 2 and 3 (Fig. 3E). Therefore, we proceeded to optimise a compact target-miRNA panel and subsequently assessed alternative normalisation schemes for the selected panel (Fig. 4).

**Fig. 4:**
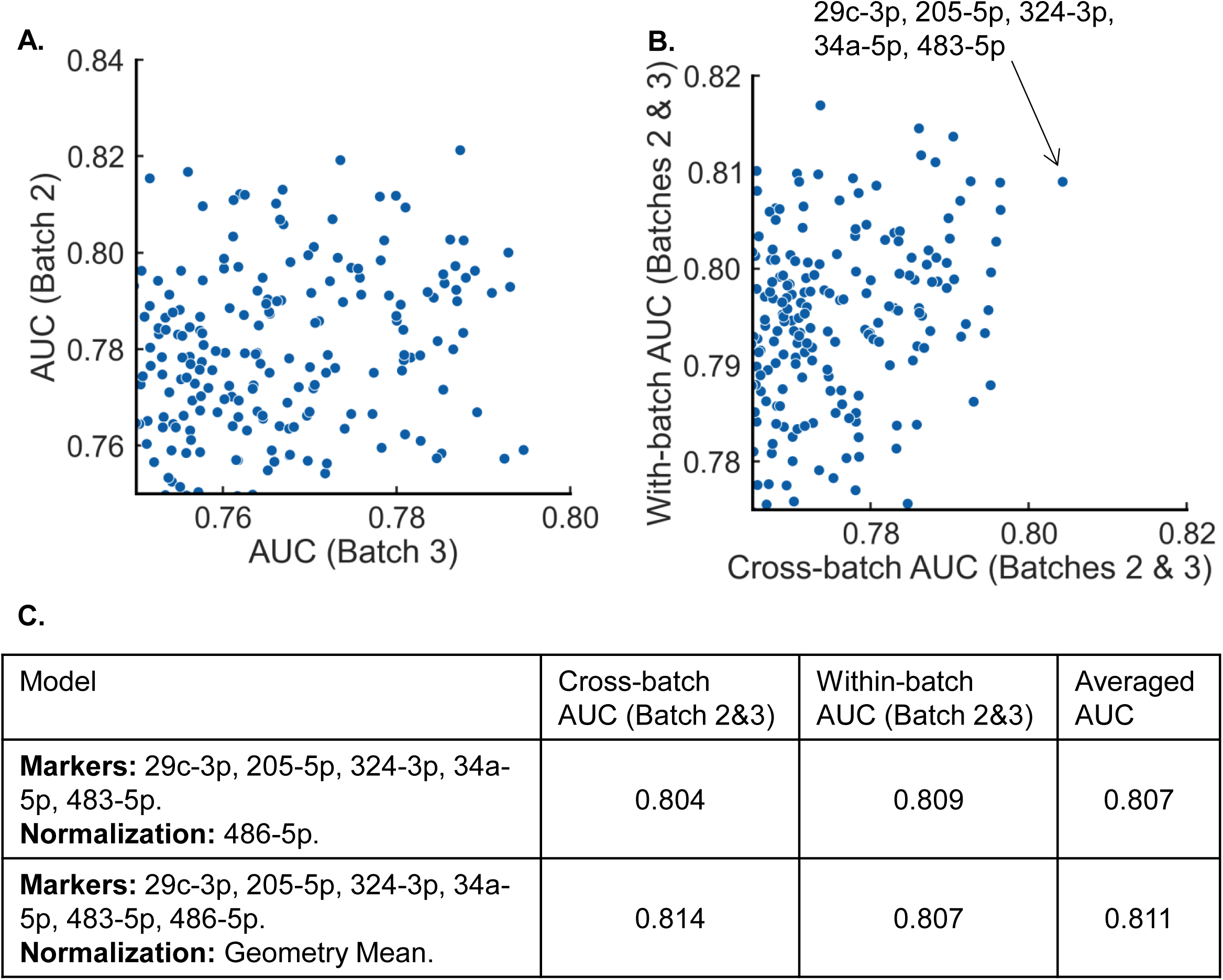
Selection of a robust compact miRNA panel. (A) Cross-batch performance of all evaluated 5 miRNA combinations derived from the 13 miRNA set. Each point represents an individual 5 miRNA panel, plotted by the AUC obtained when the models were trained on one batch and evaluated on the other (Batches 2 and 3, with reciprocal training–testing). (B) Comparison of cross-batch and within-batch AUCs for the same 5 miRNA combinations, pointing to the selected panel consisting of miR-29c-3p, miR-205-5p, miR-324-3p, miR-34a-5p, and miR-483-5p. (C) Summary of model performance under two alternative normalisation methods: normalisation to miR-486-5p and geometric-mean normalisation incorporating all measured miRNAs. Cross-batch AUC, within-batch AUC, and their average are shown. AUC, area under the curve.

### Cross-batch evaluation selected a robust miRNA model

Based on the marker-count analysis in Fig. 3E, we performed panel selection and model-configuration to define the final screening model (Fig. 4A–C). Specifically, we exhaustively evaluated all five target miRNA combinations from the 13 miRNA set, using (i) cross-batch validation between batches 2 and 3 in both directions (Fig. 4A) and (ii) within-batch validation in the dataset combining batches 2 and 3. Taking into account both cross-batch and within-batch validation, we fixed the target miRNA panel to the five miRNAs that showed consistently high AUCs across these settings: miR-29c-3p, miR-205-5p, miR-34a-5p, miR-483-5p, and miR-324-3p (Fig. 4B).

Next, we refined the model architecture to improve generalisability to unseen batches. In this refinement step, we compared two normalisation approaches while keeping the target miRNA panel and feature set constant. Under standard normalisation using miR-486-5p as an internal control, the mean AUC across metrics was 0.807 (cross-batch AUC average 0.804; within-batch AUC 0.809). When normalisation was performed using the geometric mean of the measured miRNAs, overall performance was comparable with a slightly higher cross-batch AUC average (mean AUC 0.811; cross-batch AUC average 0.814; within-batch AUC 0.807), and this refinement was carried forward for subsequent analyses (Fig. 4C). The final PC score was calculated using a fixed linear model incorporating both normalised expression values and their log10 transformed values of selected miRNAs. The model intercept was −2.31, and the corresponding coefficients were 0.0079 (miR-29c-3p), 0.214 (miR-205-5p), 1.19 (miR-324-3p), −0.0686 (miR-34a-5p), −0.366 (miR-483-5p), −1.07 (miR-486-5p), −0.688 (log10 miR-29c-3p), −1.03 (log10 miR-205-5p), 0.246 (log10 miR-324-3p), 0.112 (log10 miR-34a-5p), 0.161 (log10 miR-483-5p), and 1.02 (log10 miR-486-5p).

### Independent validation confirmed diagnostic performance and stage-wise discrimination

We next tested the final miRNA model in the independent validation batch (batch 4; n=616). Expression patterns of the miRNAs are shown in Fig. 5A (summary statistics in Supplementary Table 7). The model discriminated PC from NC with an AUC of 0.91 (Fig. 5B) and discriminated PC from NC plus other malignancies with an AUC of 0.83 (Fig. 5C), confirming generalisability in the newly collected samples. To assess the potential utility for early-stage detection, we evaluated performance by clinical stage in batch 4, using NC. The model achieved AUC values of 0.91 (I), 0.94 (II), 0.86 (III), 0.90 (IV) (Fig. 5D), indicating that the selected miRNA signature remains detectable across disease stages, including earlier-stage cases. Finally, sensitivity and specificity were evaluated at three pre-specified PC score thresholds. Thresholds selected during model training (batches 2–3) were applied to batch 4 (Fig. 5E).

**Fig. 5:**
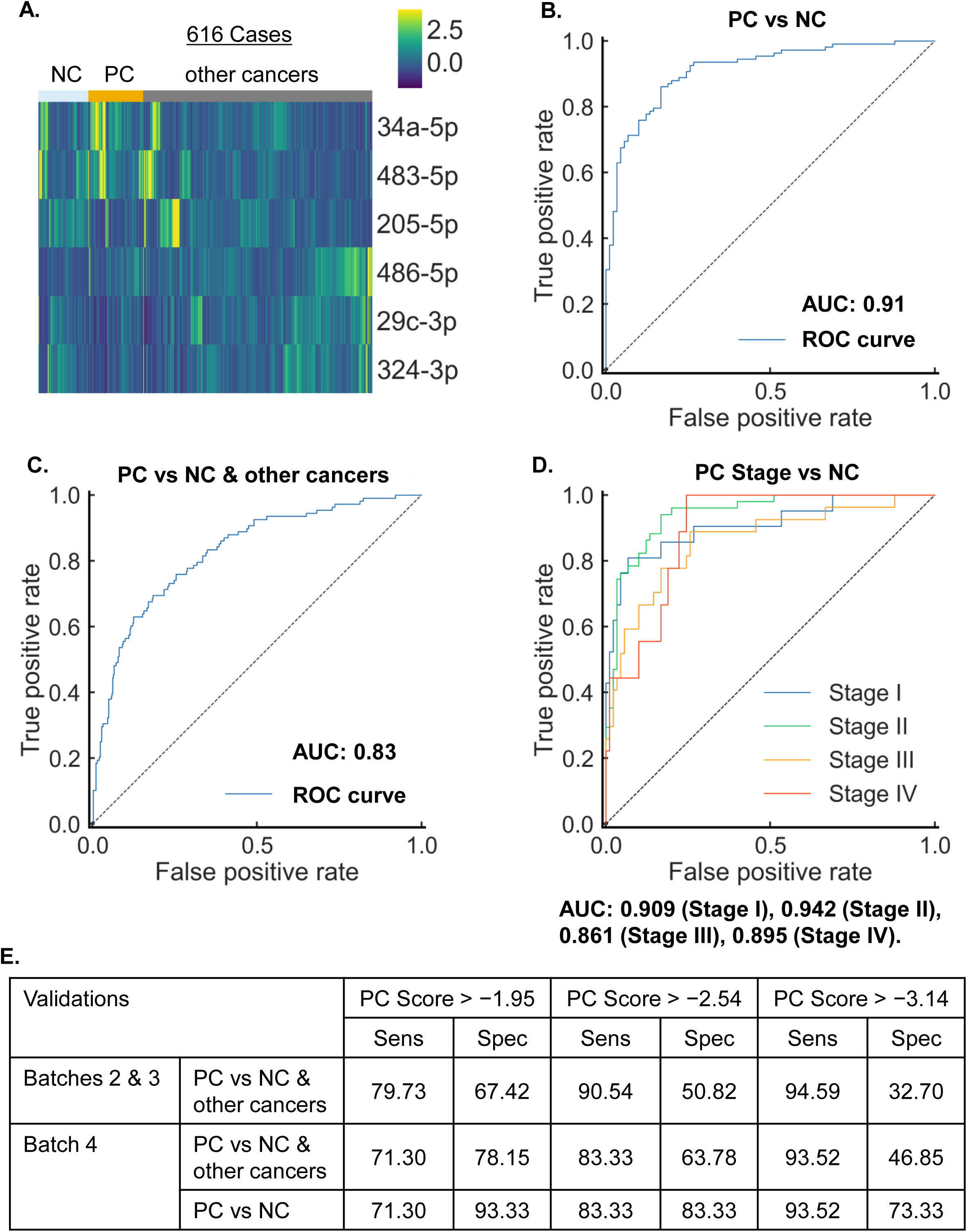
Independent validation of the final miRNA model in Batch 4. (A) Heatmap of the final miRNA panel measured in the independent validation cohort (Batch 4; n = 616; NC, PC and other cancer types), shown as scaled expression values. (B, C) ROC curves demonstrating the performance of the final model in Batch 4 for (B) PC versus NC and (C) PC versus NC plus other cancer types. (D) Stage-stratified ROC curves for discrimination of PC (stages I–IV) versus NC in Batch 4. (E) Sensitivity and specificity evaluated in Batches 2 and 3 and in Batch 4 at three pre-specified score thresholds (PC score > −1.950, > −2.54, and > −3.14). Thresholds selected during model training were applied to the independent validation cohort. AUC, area under the curve; NC, non-cancer control; PC, pancreatic cancer; ROC, receiver operating characteristic.

### Evaluation in clinical samples linked the score to surgery and recurrence

To investigate whether the selected miRNA profile also reflects disease status in a clinical setting, we reconstructed a PC status model, using only NC and PC samples to avoid potential confounding by other malignancies. This model discriminated PC status in the samples with AUC 0.94 (Fig. 6A).

**Fig. 6:**
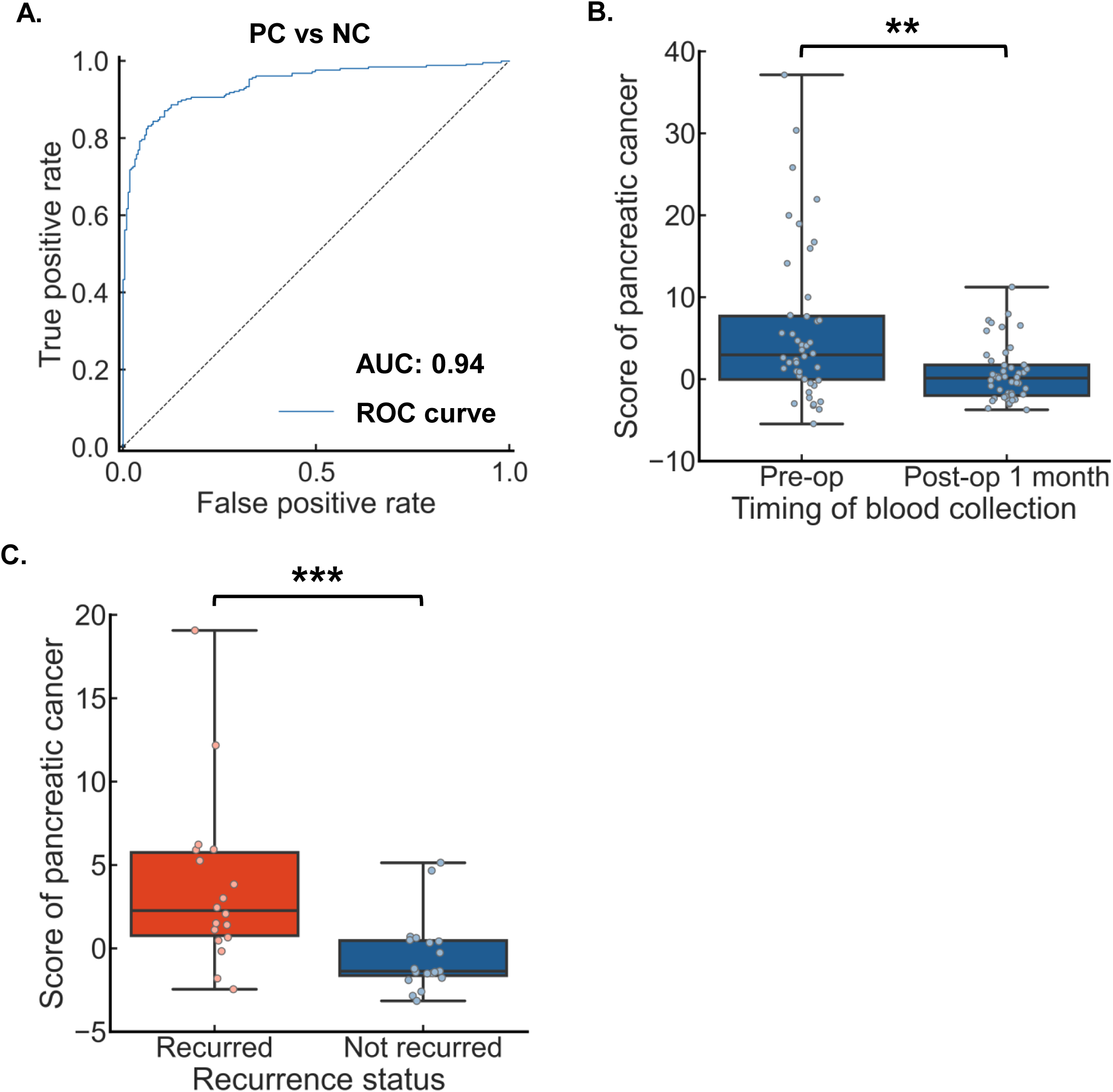
Clinical relevance of the miRNA score in perioperative monitoring and recurrence related analyses. (A) ROC curve demonstrating discrimination between PC and NC cases using the model reconstructed for clinical application, based on the selected miRNA panel. (B) Paired comparison of miRNA-derived scores between preoperative and postoperative month 1 serum samples in the clinical cohort (paired samples; n = 40). (C) Comparison of miRNA scores at postoperative month 12 between participants with recurrence and those without recurrence (recurrence group, n = 18; non-recurrence group, n = 19). In the box plots, the centre line represents the median, the box indicates the interquartile range, and the whiskers denote the data range; **p < 0.01 and ***p < 0.001. AUC, area under the curve; NC, non-cancer control; PC, pancreatic cancer; ROC, receiver operating characteristic.

We applied the model to an independent clinical cohort obtained from the National Cancer Center biobank. Serum samples were collected as paired preoperative and postoperative month 1 specimens from 40 patients (23 with recurrence and 17 without recurrence at month 12). The model-derived score significantly decreased at postoperative month 1 compared with the paired preoperative samples (Wilcoxon signed-rank test, p=0.0051; Fig. 6B).

Furthermore, serum samples were collected as postoperative month 12 specimens from 37 patients (18 recurrence, 19 non-recurrence). The score at postoperative month 12 was significantly higher in patients with recurrence than in those without recurrence (Mann–Whitney U test, p=0.00039; Fig. 6C). Taken together, these results suggest that the model-derived discrimination score tracks postoperative disease status, decreasing after surgery and increasing again in patients with recurrence at 1 year.

## Discussion

In this study, we developed a compact, qPCR-based miRNA model that discriminates PC from NC and other malignancies across multiple independent batches. The model maintained performance when other common cancers were included and retained its discriminatory ability across clinical stages, including stage I/II. We also observed clinical relevance for postoperative surveillance in an independent clinical cohort, where the miRNA-derived score decreased after surgery and was higher at postoperative month 12 in patients with recurrence within 1 year. Together, these findings indicate that the compact serum miRNA panel is compatible with routine qPCR workflows and provides evidence of cross-batch generalisability.

A major challenge in PC management is that curative-intent surgical resection is feasible for only a minority of patients, and even after surgery, recurrence is frequent and thus surveillance is essential. From a clinical perspective, serum CA19-9 remains the most widely used blood-based marker in routine practice for PC, including perioperative risk stratification and postoperative monitoring, where longitudinal kinetics can be informative^4,6,17,18^. Accordingly, miRNA-based measurements provide an early-detection perspective and complement blood-based markers, potentially improving diagnostic discrimination and enabling repeated sampling to augment established imaging- and pathology-based assessments^2–4,17–19^.

Prior work has established that circulating miRNA signatures can discriminate PC from controls using multistage discovery–validation designs, including sequencing- or array-based discovery followed by qPCR verification and risk-score approaches^11,13,20,21^. Notably, large studies have shown that blood miRNA panels can achieve AUC values of 0.9 or higher and, in some settings, provide complementary information to CA19-9^13,20,21^. However, a recurring challenge is the reproducibility of results across independent assay batches and laboratories, given that circulating miRNA measurements can be influenced by pre-analytical factors and batch effects^8,16,20^. We therefore emphasised robustness and generalisability through a staged workflow, explicit cross-batch evaluation (training on batch 2 and testing on batch 3 and vice versa), and independent validation in newly collected samples. In addition, we performed exhaustive panel selection from the refined candidate set and compared model specifications, including normalisation methods and feature formulations. Together, this framework supports the selection of a compact miRNA panel and an assay-tolerant scoring approach suitable for translation to clinically deployable qPCR-based testing.

A further practical strength is the evaluation against other malignancies rather than only healthy controls. Biomarker performance may be overestimated when controls are exclusively healthy, whereas real-world evaluation often requires distinguishing PC from other cancers and benign conditions^6,22,23^. In our multi-cancer setting, we used breast, lung, gastric, and colorectal cancers as comparators. Under these conditions, the miRNA model maintained discrimination, supporting its robustness under more clinically realistic comparisons. Stage-stratified analyses in the independent validation batch suggested that discrimination remained high even in resectable stages (stage I/II AUCs; 091/0.94). We interpret these early-stage estimates cautiously because stage-specific performance can be sensitive to batch composition and sample size. Nevertheless, the consistency across independent assay batches supports the potential of miRNA-based blood testing to provide measurable signals even in earlier-stage disease.

The selected panel also appears biologically plausible, suggesting a physiologically meaningful link to pancreatic tumourigenesis. miR-29c-3p is downregulated in PC and has been reported to suppress proliferation and invasion by targeting adhesion and signaling pathways, including ITGB1 and the MAPK1/ERK–MAPK pathway, consistent with a tumour-suppressive role^24,25^. miR-34a is a well-established tumour-suppressive miRNA that can restrain PC progression through EMT- and Notch-related signaling circuitry and has been linked to gemcitabine resistance and cancer stemness phenotypes in experimental models^26,27^. For miR-205-5p, prior studies indicate context-dependent associations with epithelial state and EMT regulators. They also suggest that combining circulating miR-205-5p with CA19-9 may improve discrimination in some cohorts, although the mechanistic role may vary across tumour contexts^28–30^. miR-324-3p has been implicated in PC malignancy via lncRNA–miRNA–mRNA axes that converge on tight-junction and pro-metastatic programmes (e.g., CLDN3), providing a plausible link to invasive dynamics^31^. Together, these miRNAs represent complementary regulatory processes implicated in PC biology.

In addition to the discrimination performance, the clinical cohort suggests that the miRNA-derived score reflects disease dynamics after resection. The significant decrease between preoperative and postoperative month 1 samples is consistent with a reduction in tumour burden after surgery, analogous to perioperative changes described for established markers such as CA19-9 as well as other circulating biomarker modalities^17–19,32^. At postoperative month 12, the score was higher in patients with recurrence than in those without recurrence, suggesting that the score may capture recurrence-associated disease status during postoperative surveillance. Although these observations do not establish a definitive prognostic biomarker, they do support the clinical relevance of the score and indicate the need for prospective studies aligned with imaging-based assessments and time-to-event endpoints.

This study has several limitations that should be kept in mind. First, the batches used for marker selection and model development were retrospective and based on sera obtained from commercial biobanks, where pre-analytical factors such as collection, processing, storage, freeze–thaw cycles, and haemolysis can influence circulating miRNA measurements. Therefore, assay standardisation and external quality control will be essential for translation^8,16,20^. Second, although we included multiple cancer types, evaluation against benign pancreatic diseases and high-risk surveillance populations will be important to define the intended use and to estimate clinically meaningful operating characteristics at prespecified thresholds^11,13,17,20^. Third, our primary performance metric was AUC, and future work should prespecify decision thresholds and report sensitivity and specificity as well as predictive values in settings with realistic disease prevalence. Finally, the clinical analyses were based on retrospective biobank specimens with limited and non-uniform sampling. The recurrence analysis focused on a single postoperative time point. Prospective studies with predefined sampling schedules and within-patient longitudinal modelling should be conducted to establish whether changes in the score track recurrence onset and to quantify clinical utility for surveillance.

In conclusion, we presented a compact miRNA qPCR model that demonstrates discrimination of PC across independent assay batches and retains its performance when other malignancies are included. The findings in a clinical cohort further suggest that the model-derived score is associated with postoperative disease status, showing a decrease after surgery and higher values in patients with recurrence, and may have clinical relevance as an adjunct measure for surveillance. Future prospective, multicentre studies with standardised workflows, predefined thresholds, and harmonised longitudinal endpoints are needed to ascertain this miRNA panel can best complement established clinical practice.

## Acknowledgments

Some of the samples and clinical information used in this study were obtained from the National Cancer Center Biobank, which is supported by National Cancer Center Research and Development Fund (29- A-1). The authors also thank the Biobank at the National Center for Geriatrics and Gerontology for providing biological resources.

## Additional information

### Data availability

The datasets generated and/or analysed during the current study are available from the corresponding author on reasonable request, subject to applicable ethical and institutional approvals.

### Ethics approval and consent to participate

The study was approved by the relevant institutional review boards, including the National Cancer Center Research Ethics Review Committee approval number 2024-023 for the clinical cohort. Written informed consent had been obtained from all participants for the use of biospecimens in research.

### Author contributions

S.Y., Y.Y. and Y.S. conceived and designed the study. HK., T.A., M.I., M.S. and M.T. performed the miRNA experiments and data acquisition. S.Y., M.T. and Y.S. performed data analysis and model development. M.E., K.K. and Y.Y. contributed to the clinical cohort, clinical data interpretation and sample-related oversight. S.Y. and Y.S. drafted the manuscript. All authors contributed to interpretation of the results, critically revised the manuscript, approved the final version, and agreed to be accountable for all aspects of the work.

## References

1. Siegel, R. L., Kratzer, T. B., Giaquinto, A. N., Sung, H. & Jemal, A. Cancer statistics, 2025. CA. Cancer J. Clin. 75, 10–45 (2025).

2. Vincent, A., Herman, J., Schulick, R., Hruban, R. H. & Goggins, M. Pancreatic cancer. Lancet 378, 607–620 (2011).

3. Kamisawa, T., Wood, L. D., Itoi, T. & Takaori, K. Pancreatic cancer. Lancet 388, 73–85 (2016).

4. Duffy, M. J. CA 19-9 as a marker for gastrointestinal cancers: a review. Ann. Clin. Biochem. 35 (Pt 3), 364–370 (1998).

5. Raghavan, S. R., Ballehaninna, U. K. & Chamberlain, R. S. The impact of perioperative blood glucose levels on pancreatic cancer prognosis and surgical outcomes: an evidence-based review. Pancreas 42, 1210–1217 (2013).

6. Steinberg, W. The clinical utility of the CA 19-9 tumor-associated antigen. Am. J. Gastroenterol. 85, 350–355 (1990).

7. Hayes, J., Peruzzi, P. P. & Lawler, S. MicroRNAs in cancer: biomarkers, functions and therapy. Trends Mol. Med. 20, 460–469 (2014).

8. He, Y. et al. Current State of Circulating MicroRNAs as Cancer Biomarkers. Clin. Chem. 61, 1138–1155 (2015).

9. Bartel, D. P. MicroRNAs: genomics, biogenesis, mechanism, and function. Cell 116, 281–297 (2004).

10. Calin, G. A. & Croce, C. M. MicroRNA signatures in human cancers. Nat. Rev. Cancer 6, 857–866 (2006).

11. Bloomston, M. et al. MicroRNA expression patterns to differentiate pancreatic adenocarcinoma from normal pancreas and chronic pancreatitis. JAMA 297, 1901–1908 (2007).

12. Mitchell, P. S. et al. Circulating microRNAs as stable blood-based markers for cancer detection. Proc. Natl. Acad. Sci. U. S. A. 105, 10513–10518 (2008).

13. Kawai, M. et al. Early detection of pancreatic cancer by comprehensive serum miRNA sequencing with automated machine learning. Br. J. Cancer 131, 1158–1168 (2024).

14. Baba, S. et al. A noninvasive urinary microRNA-based assay for the detection of pancreatic cancer from early to late stages: a case control study. EClinicalMedicine 78, 102936 (2024).

15. Matsuzaki, J., et al. Prediction of tissue-of-origin of early stage cancers using serum miRNomes. JNCI Cancer Spectr. 7, pkac080 (2023).

16. El-Daly, S. M., Gouhar, S. A. & Abd Elmageed, Z. Y. Circulating microRNAs as Reliable Tumor Biomarkers: Opportunities and Challenges Facing Clinical Application. J. Pharmacol. Exp. Ther. 384, 35–51 (2023).

17. Ferrone, C. R. et al. Perioperative CA19-9 levels can predict stage and survival in patients with resectable pancreatic adenocarcinoma. J. Clin. Oncol. Off. J. Am. Soc. Clin. Oncol. 24, 2897–2902 (2006).

18. Humphris, J. L. et al. The prognostic and predictive value of serum CA19.9 in pancreatic cancer. Ann. Oncol. Off. J. Eur. Soc. Med. Oncol. 23, 1713–1722 (2012).

19. Azizian, A. et al. CA19-9 for detecting recurrence of pancreatic cancer. Sci. Rep. 10, 1332 (2020).

20. Liu, R. et al. Serum microRNA expression profile as a biomarker in the diagnosis and prognosis of pancreatic cancer. Clin. Chem. 58, 610–618 (2012).

21. Schultz, N. A. et al. MicroRNA biomarkers in whole blood for detection of pancreatic cancer. JAMA 311, 392–404 (2014).

22. Ransohoff, D. F. & Feinstein, A. R. Problems of spectrum and bias in evaluating the efficacy of diagnostic tests. N. Engl. J. Med. 299, 926–930 (1978).

23. Melo, S. A. et al. Glypican-1 identifies cancer exosomes and detects early pancreatic cancer. Nature 523, 177–182 (2015).

24. Lu, Y. et al. MiR-29c inhibits cell growth, invasion, and migration of pancreatic cancer by targeting ITGB1. OncoTargets Ther. 9, 99–109 (2016).

25. Si, H., Zhang, N., Shi, C., Luo, Z. & Hou, S. Tumor-suppressive miR-29c binds to MAPK1 inhibiting the ERK/MAPK pathway in pancreatic cancer. Clin. Transl. Oncol. Off. Publ. Fed. Span. Oncol. Soc. Natl. Cancer Inst. Mex. 25, 803–816 (2023).

26. Tang, Y., Tang, Y. & Cheng, Y.-S. miR-34a inhibits pancreatic cancer progression through Snail1-mediated epithelial-mesenchymal transition and the Notch signaling pathway. Sci. Rep. 7, 38232 (2017).

27. Pan, Y. et al. MicroRNA-34a Alleviates Gemcitabine Resistance in Pancreatic Cancer by Repression of Cancer Stem Cell Renewal. Pancreas 50, 1260–1266 (2021).

28. Wang, Y. et al. Extracellular vesicles-miR-205-5p inhibits lymphatic metastasis in pancreatic cancer through diffusely downregulating VEGFA. J. Cancer 16, 2197–2211 (2025).

29. Gregory, P. A. et al. The miR-200 family and miR-205 regulate epithelial to mesenchymal transition by targeting ZEB1 and SIP1. Nat. Cell Biol. 10, 593–601 (2008).

30. Michael Traeger, M., et al. The ambiguous role of microRNA-205 and its clinical potential in pancreatic ductal adenocarcinoma. J. Cancer Res. Clin. Oncol. 144, 2419–2431 (2018).

31. Jiang, F., Li, S., Wang, X., Deng, Y. & Peng, S. DPP10-AS1-Mediated Downregulation of MicroRNA-324-3p Is Conducive to the Malignancy of Pancreatic Cancer by Enhancing CLDN3 Expression. Pancreas 51, 1201–1210 (2022).

32. Sausen, M. et al. Clinical implications of genomic alterations in the tumour and circulation of pancreatic cancer patients. Nat. Commun. 6, 7686 (2015).

