## Supplemental File for "Compact serum miRNA qPCR model for pancreatic cancer discrimination with independent and clinical validation"

**Supplementary Table 1: Sample information for model development and validation cohorts.**

| | Types of cancer | Stage | | | | | Sex | | Age | | Total |
|---|---|---|---|---|---|---|---|---|---|---|---|
| | | 0 | I | II | III | IV | M | F | ≤60 | ≥61 | |
| **Batch 1** | **Non-cancer** | | | | | | 18 | 6 | 17 | 7 | 24 |
| | **Pancreatic cancer** | | 3 | 18 | 2 | 1 | 13 | 11 | 13 | 11 | 24 |
| | **Breast cancer** | 2 | 9 | 13 | | | | 24 | 17 | 7 | 24 |
| **Batch 2** | **Non-cancer** | | | | | | 44 | 37 | 60 | 21 | 81 |
| | **Pancreatic cancer** | | 19 | 48 | 10 | 5 | 34 | 48 | 30 | 52 | 82 |
| | **Breast cancer** | 6 | 28 | 55 | 10 | | | 99 | 63 | 36 | 99 |
| | **Lung cancer** | 1 | 38 | 36 | 21 | 2 | 69 | 29 | 60 | 38 | 98 |
| | **Gastric cancer** | 4 | 30 | 36 | 26 | 1 | 53 | 44 | 62 | 35 | 97 |
| | **Colorectal cancer** | 5 | 29 | 30 | 31 | | 51 | 44 | 54 | 41 | 95 |
| **Batch 3** | **Non-cancer** | | | | | | 25 | 37 | 32 | 30 | 62 |
| | **Pancreatic cancer** | | 14 | 29 | 6 | 17 | 32 | 34 | 33 | 33 | 66 |
| | **Breast cancer** | 1 | 26 | 36 | 5 | 1 | | 69 | 47 | 22 | 69 |
| | **Lung cancer** | | 24 | 16 | 4 | 19 | 41 | 22 | 25 | 38 | 63 |
| | **Gastric cancer** | 2 | 27 | 24 | 11 | | 38 | 26 | 25 | 39 | 64 |
| | **Colorectal cancer** | 3 | 19 | 14 | 23 | 8 | 39 | 28 | 36 | 31 | 67 |
| **Batch 4** | **Non-cancer** | | | | | | 43 | 47 | 57 | 33 | 90 |
| | **Pancreatic cancer** | | 21 | 51 | 27 | 9 | 42 | 66 | 40 | 68 | 108 |
| | **Breast cancer** | 7 | 42 | 46 | 10 | | | 105 | 69 | 36 | 105 |
| | **Lung cancer** | 1 | 38 | 19 | 35 | 11 | 71 | 33 | 37 | 67 | 104 |
| | **Gastric cancer** | | 31 | 40 | 22 | 8 | 55 | 46 | 42 | 59 | 101 |
| | **Colorectal cancer** | 2 | 24 | 42 | 35 | 5 | 57 | 51 | 47 | 61 | 108 |

**Supplementary Table 2: Sample information for the clinical cohort.**

| | Types of cancer | Stage | | | | | Sex | | Age | | Total |
|---|---|---|---|---|---|---|---|---|---|---|---|
| | | 0 | I | II | III | IV | M | F | ≤60 | ≥61 | |
| Clinical cohort | Not-recurred | | 15 | 5 | | | 11 | 9 | 5 | 15 | 20 |
| | Recurred | 1 | 15 | 13 | 2 | 1 | 16 | 16 | 5 | 27 | 32 |

**Supplementary Table 3-1: TaqMan probe information and target sequences for qPCR-based miRNA analysis.**

| Marker | Probe ID | miRBase ID | Mature sequence |
|---|---|---|---|
| hsa-miR-107 | 478254_mir | MIMAT0000104 | AGCAGCAUUGUACAGGGCUAUCA |
| hsa-miR-326 | 478027_mir | MIMAT0000756 | CCUCUGGGCCCUUCCUCCAG |
| hsa-miR-484 | 478308_mir | MIMAT0002174 | UCAGGCUCAGUCCCCUCCCGAU |
| hsa-miR-559 | 479045_mir | MIMAT0003223 | UAAAGUAAAUAUGCACCAAAA |
| hsa-miR-606 | 479087_mir | MIMAT0003274 | AAACUACUGAAAAUCAAAGAU |
| hsa-miR-665 | 479150_mir | MIMAT0004952 | ACCAGGAGGCUGAGGCCCCU |
| hsa-miR-941 | 479217_mir | MIMAT0004984 | CACCCGGCUGUGUGCACAUGUGC |
| hsa-miR-1246 | 483023_mir | MIMAT0005898 | AAUGGAUUUUUGGAGCAGG |
| hsa-miR-3168 | 480795_mir | MIMAT0015043 | GAGUUCUACAGUCAGAC |
| hsa-miR-3178 | 483330_mir | MIMAT0015055 | GGGGCGCGGCCGGAUCG |
| hsa-miR-3182 | 480797_mir | MIMAT0015062 | GCUUCUGUAGUGUAGUC |
| hsa-miR-3615 | 478837_mir | MIMAT0017994 | UCUCUCGGCUCCUCGCGGCUC |
| hsa-miR-4300 | 480810_mir | MIMAT0016853 | UGGGAGCUGGACUACUUC |
| hsa-miR-4302 | 478099_mir | MIMAT0016855 | CCAGUGUGGCUCAGCGAG |
| hsa-miR-4466 | 483160_mir | MIMAT0018993 | GGGUGCGGGCCGGCGGGG |
| hsa-miR-4770 | 478120_mir | MIMAT0019924 | UGAGAUGACACUGUAGCU |
| hsa-miR-101-3p | 477863_mir | MIMAT0000099 | UACAGUACUGUGAUAACUGAA |
| hsa-miR-103a-3p | 478253_mir | MIMAT0000101 | AGCAGCAUUGUACAGGGCUAUGA |
| hsa-miR-106b-5p | 478412_mir | MIMAT0000680 | UAAAGUGCUGACAGUGCAGAU |
| hsa-miR-10b-5p | 478494_mir | MIMAT0000254 | UACCCUGUAGAACCGAAUUUGUG |
| hsa-miR-122-5p | 477855_mir | MIMAT0000421 | UGGAGUGUGACAAUGGUGUUUG |
| hsa-miR-1229-3p | 478645_mir | MIMAT0005584 | CUCUCACCACUGCCCUCCCACAG |
| hsa-miR-1249-3p | 478654_mir | MIMAT0005901 | ACGCCCUUCCCCCCCUUCUUCA |
| hsa-miR-125a-5p | 477884_mir | MIMAT0000443 | UCCCUGAGACCCUUUAACCUGUGA |
| hsa-miR-125b-5p | 477885_mir | MIMAT0000423 | UCCCUGAGACCCUAACUUGUGA |
| hsa-miR-126-3p | 477887_mir | MIMAT0000445 | UCGUACCGUGAGUAAUAAUGCG |
| hsa-miR-127-3p | 477889_mir | MIMAT0000446 | UCGGAUCCGUCUGAGCUUGGCU |
| hsa-miR-128-3p | 477892_mir | MIMAT0000424 | UCACAGUGAACCGGUCUCUUU |
| hsa-miR-1296-5p | 479451_mir | MIMAT0005794 | UUAGGGCCCUGGCUCCAUCUCC |
| hsa-miR-1307-3p | 483036_mir | MIMAT0005951 | ACUCGGCGUGGCGUCGGUCGUG |
| hsa-miR-132-3p | 477900_mir | MIMAT0000426 | UAACAGUCUACAGCCAUGGUCG |
| hsa-miR-133a-3p | 478511_mir | MIMAT0000427 | UUUGGUCCCCUUCAACCAGCUG |
| hsa-miR-1-3p | 477820_mir | MIMAT0000416 | UGGAAUGUAAAGAAGUAUGUAU |
| hsa-miR-140-3p | 477908_mir | MIMAT0004597 | UACCACAGGGUAGAACCACGG |
| hsa-miR-142-3p | 477910_mir | MIMAT0000434 | UGUAGUGUUUCCUACUUUAUGGA |
| hsa-miR-142-5p | 477911_mir | MIMAT0000433 | CAUAAAGUAGAAAGCACUACU |
| hsa-miR-143-3p | 477912_mir | MIMAT0000435 | UGAGAUGAAGCACUGUAGCUC |
| hsa-miR-148a-3p | 477814_mir | MIMAT0000243 | UCAGUGCACUACAGAACUUUGU |
| hsa-miR-148b-3p | 477824_mir | MIMAT0000759 | UCAGUGCAUCACAGAACUUUGU |
| hsa-miR-150-5p | 477918_mir | MIMAT0000451 | UCUCCCAACCCUUGUACCAGUG |
| hsa-miR-151a-3p | 477919_mir | MIMAT0000757 | CUAGACUGAAGCUCCUUGAGG |
| hsa-miR-155-5p | 483064_mir | MIMAT0000646 | UUAAUGCUAAUCGUGAUAGGGGUU |
| hsa-miR-15a-5p | 477858_mir | MIMAT0000068 | UAGCAGCACAUAAUGGUUUGUG |
| hsa-miR-15b-5p | 478313_mir | MIMAT0000417 | UAGCAGCACAUCAUGGUUUACA |
| hsa-miR-16-2-3p | 477931_mir | MIMAT0004518 | CCAAUAUUACUGUGCUGCUUUA |
| hsa-miR-16-5p | 477860_mir | MIMAT0000069 | UAGCAGCACGUAAAUAUUGGCG |
| hsa-miR-17-5p | 478447_mir | MIMAT0000070 | CAAAGUGCUUACAGUGCAGGUAG |
| hsa-miR-181a-5p | 477857_mir | MIMAT0000256 | AACAUUCAACGCUGUCGGUGAGU |
| hsa-miR-181b-5p | 478583_mir | MIMAT0000257 | AACAUUCAUUGCUGUCGGUGGGU |
| hsa-miR-186-5p | 477940_mir | MIMAT0000456 | CAAAGAAUUCUCCUUUUGGGCU |
| hsa-miR-18a-5p | 478551_mir | MIMAT0000072 | UAAGGUGCAUCUAGUGCAGAUAG |
| hsa-miR-190a-5p | 478358_mir | MIMAT0000458 | UGAUAUGUUUGAUAUAUUAGGU |
| hsa-miR-191-5p | 477952_mir | MIMAT0000440 | CAACGGAAUCCCAAAAGCAGCUG |
| hsa-miR-192-5p | 478262_mir | MIMAT0000222 | CUGACCUAUGAAUUGACAGCC |

**Supplementary Table 3-2: TaqMan probe information and target sequences for qPCR-based miRNA analysis.**

| Marker | Probe ID | miRBase ID | Mature sequence |
|---|---|---|---|
| hsa-miR-193a-5p | 477954_mir | MIMAT0004614 | UGGGUCUUUGCGGGCGAGAUGA |
| hsa-miR-194-5p | 477956_mir | MIMAT0000460 | UGUAACAGCAACUCCAUGUGGA |
| hsa-miR-196a-5p | 478230_mir | MIMAT0000226 | UAGGUAGUUUCAUGUUGUUGGG |
| hsa-miR-197-3p | 477959_mir | MIMAT0000227 | UUCACCACCUUCUCCACCCAGC |
| hsa-miR-199a-3p | 477961_mir | MIMAT0000232 | ACAGUAGUCUGCACAUUGGUUA |
| hsa-miR-19b-3p | 478264_mir | MIMAT0000074 | UGUGCAAAUCCAUGCAAAACUGA |
| hsa-miR-205-5p | 477967_mir | MIMAT0000266 | UCCUUCAUUCCACCGGAGUCUG |
| hsa-miR-20a-5p | 478586_mir | MIMAT0000075 | UAAAGUGCUUAUAGUGCAGGUAG |
| hsa-miR-215-5p | 478516_mir | MIMAT0000272 | AUGACCUAUGAAUUGACAGAC |
| hsa-miR-21-5p | 477975_mir | MIMAT0000076 | UAGCUUAUCAGACUGAUGUUGA |
| hsa-miR-221-3p | 477981_mir | MIMAT0000278 | AGCUACAUUGUCUGCUGGGUUUC |
| hsa-miR-222-3p | 477982_mir | MIMAT0000279 | AGCUACAUCUGGCUACUGGGU |
| hsa-miR-223-3p | 477983_mir | MIMAT0000280 | UGUCAGUUUGUCAAAUACCCCA |
| hsa-miR-223-5p | 477984_mir | MIMAT0004570 | CGUGUAUUUGACAAGCUGAGUU |
| hsa-miR-22-3p | 477985_mir | MIMAT0000077 | AAGCUGCCAGUUGAAGAACUGU |
| hsa-miR-22-5p | 477987_mir | MIMAT0004495 | AGUUCUUCAGUGGCAAGCUUUA |
| hsa-miR-23a-3p | 478532_mir | MIMAT0000078 | AUCACAUUGCCAGGGAUUUCC |
| hsa-miR-23b-3p | 483150_mir | MIMAT0000418 | AUCACAUUGCCAGGGAUUACCAC |
| hsa-miR-24-3p | 477992_mir | MIMAT0000080 | UGGCUCAGUUCAGCAGGAACAG |
| hsa-miR-26b-5p | 478418_mir | MIMAT0000083 | UUCAAGUAAUUCAGGAUAGGU |
| hsa-miR-27a-3p | 478384_mir | MIMAT0000084 | UUCACAGUGGCUAAGUUCCGC |
| hsa-miR-27b-3p | 478270_mir | MIMAT0000419 | UUCACAGUGGCUAAGUUCUGC |
| hsa-miR-27b-5p | 478789_mir | MIMAT0004588 | AGAGCUUAGCUGAUUGGUGAAC |
| hsa-miR-28-3p | 477999_mir | MIMAT0004502 | CACUAGAUUGUGAGCUCCUGGA |
| hsa-miR-29a-3p | 478587_mir | MIMAT0000086 | UAGCACCAUCUGAAAUCGGUUA |
| hsa-miR-29c-3p | 479229_mir | MIMAT0000681 | UAGCACCAUUUGAAAUCGGUUA |
| hsa-miR-30d-5p | 478606_mir | MIMAT0000245 | UGUAAACAUCCCCGACUGGAAG |
| hsa-miR-30e-5p | 479235_mir | MIMAT0000692 | UGUAAACAUCCUUGACUGGAAG |
| hsa-miR-3155a | 480794_mir | MIMAT0015029 | CCAGGCUCUGCAGUGGGAACU |
| hsa-miR-31-5p | 478015_mir | MIMAT0000089 | AGGCAAGAUGCUGGCAUAGCU |
| hsa-miR-320a-3p | 478594_mir | MIMAT0000510 | AAAAGCUGGGUUGAGAGGGCGA |
| hsa-miR-320b | 478588_mir | MIMAT0005792 | AAAAGCUGGGUUGAGAGGGCAA |
| hsa-miR-324-3p | 483109_mir | MIMAT0000762 | CCCACUGCCCCAGGUGCUGCUGG |
| hsa-miR-324-5p | 483066_mir | MIMAT0000761 | CGCAUCCCCUAGGGCAUUGGUG |
| hsa-miR-32-5p | 478026_mir | MIMAT0000090 | UAUUGCACAUUACUAAGUUGCA |
| hsa-miR-328-3p | 478028_mir | MIMAT0000752 | CUGGCCCUCUCUGCCCUUCCGU |
| hsa-miR-329-5p | 478829_mir | MIMAT0026555 | GAGGUUUUCUGGGUUUCUGUUUC |
| hsa-miR-338-5p | 478038_mir | MIMAT0004701 | AACAAUAUCCUGGUGCUGAGUG |
| hsa-miR-339-5p | 478040_mir | MIMAT0000764 | UCCCUGUCCUCCAGGAGCUCACG |
| hsa-miR-342-3p | 478043_mir | MIMAT0000753 | UCUCACACAGAAAUCGCACCCGU |
| hsa-miR-34a-5p | 478048_mir | MIMAT0000255 | UGGCAGUGUCUUAGCUGGUUGU |
| hsa-miR-3613-3p | 478434_mir | MIMAT0017991 | ACAAAAAAAAAAGCCCAACCCUUC |
| hsa-miR-361-5p | 478056_mir | MIMAT0000703 | UUAUCAGAAUCUCCAGGGGUAC |
| hsa-miR-375-3p | 478074_mir | MIMAT0000728 | UUUGUUCGUUCGGCUCGCGUGA |
| hsa-miR-378a-3p | 478349_mir | MIMAT0000732 | ACUGGACUUGGAGUCAGAAGGC |
| hsa-miR-382-5p | 478078_mir | MIMAT0000737 | GAAGUUGUUCGUGGUGGAUUCG |
| hsa-miR-409-3p | 478084_mir | MIMAT0001639 | GAAUGUUGCUCGGUGAACCCCU |
| hsa-miR-423-3p | 478327_mir | MIMAT0001340 | AGCUCGGUCUGAGGCCCCUCAGU |
| hsa-miR-423-5p | 478090_mir | MIMAT0004748 | UGAGGGGCAGAGAGCGAGACUUU |
| hsa-miR-424-3p | 478091_mir | MIMAT0004749 | CAAAACGUGAGGCGCUGCUAU |
| hsa-miR-425-3p | 478093_mir | MIMAT0001343 | AUCGGGAAUGUCGUGUCCGCCC |
| hsa-miR-433-3p | 478102_mir | MIMAT0001627 | AUCAUGAUGGGCUCCUCGGUGU |
| hsa-miR-4433a-3p | 478897_mir | MIMAT0018949 | ACAGGAGUGGGGGUGGGACAU |
| hsa-miR-4446-3p | 479812_mir | MIMAT0018965 | CAGGGCUGGCAGUGACAUGGGU |

**Supplementary Table 3-3: TaqMan probe information and target sequences for qPCR-based miRNA analysis.**

| Marker | Probe ID | miRBase ID | Mature sequence |
|---|---|---|---|
| hsa-miR-451a | 478107_mir | MIMAT0001631 | AAACCGUUACCAUUACUGAGUU |
| hsa-miR-4732-5p | 478119_mir | MIMAT0019855 | UGUAGAGCAGGGAGCAGGAAGCU |
| hsa-miR-483-3p | 478122_mir | MIMAT0002173 | UCACUCCUCUCCUCCCGUCUU |
| hsa-miR-483-5p | 478432_mir | MIMAT0004761 | AAGACGGGAGGAAAGAAGGGAG |
| hsa-miR-485-3p | 478125_mir | MIMAT0002176 | GUCAUACACGGCUCUCCUCUCU |
| hsa-miR-485-5p | 478126_mir | MIMAT0002175 | AGAGGCUGGCCGUGAUGAAUUC |
| hsa-miR-486-3p | 478422_mir | MIMAT0004762 | CGGGGCAGCUCAGUACAGGAU |
| hsa-miR-493-3p | 478134_mir | MIMAT0003161 | UGAAGGUCUACUGUGUGCCAGG |
| hsa-miR-5007-3p | 480727_mir | MIMAT0021036 | AUCAUAUGAACCAAACUCUAAU |
| hsa-miR-505-5p | 478957_mir | MIMAT0004776 | GGGAGCCAGGAAGUAUUGAUGU |
| hsa-miR-513-5p | 479297_mir | MIMAT0005788 | UUCACAAGGAGGUGUCAUUUAU |
| hsa-miR-520e-3p | 478498_mir | MIMAT0002825 | AAAGUGCUUCCUUUUUGAGGG |
| hsa-miR-532-5p | 478151_mir | MIMAT0002888 | CAUGCCUUGAGUGUAGGACCGU |
| hsa-miR-542-3p | 478153_mir | MIMAT0003389 | UGUGACAGAUUGAUAACUGAAA |
| hsa-miR-574-5p | 479357_mir | MIMAT0004795 | UGAGUGUGUGUGUGUGAGUGUGU |
| hsa-miR-576-3p | 478164_mir | MIMAT0004796 | AAGAUGUGGAAAAAUUGGAAUC |
| hsa-miR-584-5p | 478167_mir | MIMAT0003249 | UUAUGGUUUGCCUGGGACUGAG |
| hsa-miR-625-3p | 478179_mir | MIMAT0004808 | GACUAUAGAACUUUCCCCCUCA |
| hsa-miR-629-5p | 478183_mir | MIMAT0004810 | UGGGUUUACGUUGGGAGAACU |
| hsa-miR-652-3p | 478189_mir | MIMAT0003322 | AAUGGCGCCACUAGGGUUGUG |
| hsa-miR-660-5p | 478192_mir | MIMAT0003338 | UACCCAUUGCAUAUCGGAGUUG |
| hsa-miR-664a-3p | 478193_mir | MIMAT0005949 | UAUUCAUUUAUCCCCAGCCUACA |
| hsa-miR-6756-5p | 480284_mir | MIMAT0027412 | AGGGUGGGGCUGGAGGUGGGGCU |
| hsa-miR-6859-5p | 480474_mir | MIMAT0027618 | GAGAGGAACAUGGGCUCAGGACA |
| hsa-miR-6886-3p | 480521_mir | MIMAT0027673 | UGCCCUUCUCUCCUCCUGCCU |
| hsa-miR-744-5p | 478200_mir | MIMAT0004945 | UGCGGGGCUAGGGCUAACAGCA |
| hsa-miR-758-3p | 479166_mir | MIMAT0003879 | UUUGUGACCUGGUCCACUAACC |
| hsa-miR-769-5p | 478203_mir | MIMAT0003886 | UGAGACCUCUGGGUUCUGAGCU |
| hsa-let-7a-5p | 478575_mir | MIMAT0000062 | UGAGGUAGUAGGUUGUAUAGUU |
| hsa-let-7b-3p | 478221_mir | MIMAT0004482 | CUAUACAACCUACUGCCUUCCC |
| hsa-let-7b-5p | 478576_mir | MIMAT0000063 | UGAGGUAGUAGGUUGUGUGGUU |
| hsa-let-7c-5p | 478577_mir | MIMAT0000064 | UGAGGUAGUAGGUUGUAUGGUU |
| hsa-let-7d-3p | 477848_mir | MIMAT0004484 | CUAUACGACCUGCUGCCUUUCU |
| hsa-let-7e-5p | 478579_mir | MIMAT0000066 | UGAGGUAGGAGGUUGUAUAGUU |
| hsa-let-7f-5p | 478578_mir | MIMAT0000067 | UGAGGUAGUAGAUUGUAUAGUU |
| hsa-let-7g-5p | 478580_mir | MIMAT0000414 | UGAGGUAGUAGUUUGUACAGUU |
| hsa-let-7i-5p | 478375_mir | MIMAT0000415 | UGAGGUAGUAGUUUGUGCUGUU |
| hsa-miR-885-5p | 478207_mir | MIMAT0004947 | UCCAUUACACUACCCUGCCUCU |
| hsa-miR-92b-3p | 477823_mir | MIMAT0003218 | UAUUGCACUCGUCCCGGCCUCC |
| hsa-miR-93-3p | 478209_mir | MIMAT0004509 | ACUGCUGAGCUAGCACUUCCCG |
| hsa-miR-9-5p | 478214_mir | MIMAT0000441 | UCUUUGGUUAUCUAGCUGUAUGA |
| hsa-miR-98-5p | 478590_mir | MIMAT0000096 | UGAGGUAGUAAGUUGUAUUGUU |
| hsa-miR-99a-5p | 478519_mir | MIMAT0000097 | AACCCGUAGAUCCGAUCUUGUG |
| hsa-miR-99b-5p | 478343_mir | MIMAT0000689 | CACCCGUAGAACCGACCUUGCG |
| hsa-miR-206 | 477968_mir | MIMAT0000462 | UGGAAUGUAAGGAAGUGUGUGG |
| hsa-miR-146a-5p | 478399_mir | MIMAT0000449 | UGAGAACUGAAUUCCAUGGGUU |
| hsa-miR-196b-5p | 478585_mir | MIMAT0001080 | UAGGUAGUUUCCUGUUGUUGGG |
| hsa-miR-2276-3p | 477988_mir | MIMAT0011775 | UCUGCAAGUGUCAGAGGCGAGG |
| hsa-miR-25-3p | 477994_mir | MIMAT0000081 | CAUUGCACUUGUCUCGGUCUGA |
| hsa-miR-3151-5p | 478811_mir | MIMAT0015024 | GGUGGGGCAAUGGGAUCAGGU |
| hsa-miR-425-5p | 478094_mir | MIMAT0003393 | AAUGACACGAUCACUCCCGUUGA |
| hsa-miR-4433b-5p | 479803_mir | MIMAT0030413 | AUGUCCCACCCCCACUCCUGU |
| hsa-miR-574-3p | 478163_mir | MIMAT0003239 | CACGCUCAUGCACACACCCACA |
| hsa-miR-93-5p | 478210_mir | MIMAT0000093 | CAAAGUGCUGUUCGUGCAGGUAG |

**Supplementary Table 3-4: TaqMan probe information and target sequences for qPCR-based miRNA analysis.**

| Marker | Probe ID | miRBase ID | Mature sequence |
|---|---|---|---|
| hsa-miR-432-5p | 478101_mir | MIMAT0002814 | UCUUGGAGUAGGUCAUUGGGUGG |
| hsa-miR-6872-5p | 480495_mir | MIMAT0027644 | UCUCGCAUCAGGAGGCAAGG |
| hsa-miR-616-3p | 478177_mir | MIMAT0004805 | AGUCAUUGGAGGGUUUGAGCAG |
| hsa-miR-2052 | 480639_mir | MIMAT0009977 | UGUUUUGAUAACAGUAAUGU |
| hsa-miR-193b-5p | 478742_mir | MIMAT0004767 | CGGGGUUUUGAGGGCGAGAUGA |
| hsa-miR-3613-5p | 479424_mir | MIMAT0017990 | UGUUGUACUUUUUUUUUUGUUC |
| hsa-miR-6830-3p | 480423_mir | MIMAT0027561 | UGUCUUUCUUCUCUCCCUUGCAG |
| hsa-miR-148a-5p | 478718_mir | MIMAT0004549 | AAAGUUCUGAGACACUCCGACU |
| hsa-miR-6797-3p | 480360_mir | MIMAT0027495 | UGCAUGACCCUUCCCUCCCCAC |
| hsa-miR-1298-5p | 479452_mir | MIMAT0005800 | UUCAUUCGGCUGUCCAGAUGUA |
| hsa-miR-876-3p | 479186_mir | MIMAT0004925 | UGGUGGUUUACAAAGUAAUUCA |
| hsa-miR-1910-5p | 477949_mir | MIMAT0007884 | CCAGUCCUGUGCCUGCCGCCU |
| hsa-miR-509-3-5p | 478963_mir | MIMAT0004975 | UACUGCAGACGUGGCAAUCAUG |
| hsa-miR-3074-5p | 479606_mir | MIMAT0019208 | GUUCCUGCUGAACUGAGCCAG |
| hsa-miR-6131 | 480187_mir | MIMAT0024615 | GGCUGGUCAGAUGGGAGUG |
| hsa-miR-3150b-3p | 479635_mir | MIMAT0018194 | UGAGGAGAUCGUCGAGGUUGG |
| hsa-miR-3927-5p | 479748_mir | MIMAT0022970 | GCCUAUCACAUAUCUGCCUGU |
| hsa-miR-4657 | 479891_mir | MIMAT0019724 | AAUGUGGAAGUGGUCUGAGGCAU |
| hsa-miR-4699-5p | 479939_mir | MIMAT0019794 | AGAAGAUUGCAGAGUAAGUUCC |
| hsa-miR-5585-3p | 480128_mir | MIMAT0022286 | CUGAAUAGCUGGGACUACAGGU |
| hsa-miR-6873-3p | 480496_mir | MIMAT0027647 | UUCUCUCUGUCUUUCUCUCUCAG |
| hsa-miR-626 | 479110_mir | MIMAT0003295 | AGCUGUCUGAAAAUGUCUU |
| hsa-miR-1276 | 478680_mir | MIMAT0005930 | UAAAGAGCCCUGUGGAGACA |
| hsa-miR-3163 | 479648_mir | MIMAT0015037 | UAUAAAAUGAGGGCAGUAAGAC |
| hsa-miR-3662 | 478844_mir | MIMAT0018083 | GAAAAUGAUGAGUAGUGACUGAUG |
| hsa-miR-3666 | 479702_mir | MIMAT0018088 | CAGUGCAAGUGUAGAUGCCGA |
| hsa-miR-3941 | 478871_mir | MIMAT0018357 | UUACACACAACUGAGGAUCAUA |
| hsa-miR-4254 | 478875_mir | MIMAT0016884 | GCCUGGAGCUACUCCACCAUCUC |
| hsa-miR-4322 | 478891_mir | MIMAT0016873 | CUGUGGGCUCAGCGCGUGGGG |
| hsa-miR-4440 | 479809_mir | MIMAT0018958 | UGUCGUGGGGCUUGCUGGCUUG |
| hsa-miR-4501 | 479837_mir | MIMAT0019037 | UAUGUGACCUCGGAUGAAUCA |
| hsa-miR-4502 | 479838_mir | MIMAT0019038 | GCUGAUGAUGAUGGUGCUGAAG |
| hsa-miR-4540 | 479863_mir | MIMAT0019083 | UUAGUCCUGCCUGUAGGUUUA |
| hsa-miR-4765 | 480713_mir | MIMAT0019916 | UGAGUGAUUGAUAGCUAUGUUC |
| hsa-miR-4771 | 480714_mir | MIMAT0019925 | AGCAGACUUGACCUACAAUUA |
| hsa-miR-4780 | 480041_mir | MIMAT0019939 | ACCCUUGAGCCUGAUCCCUAGC |
| hsa-miR-4803 | 480724_mir | MIMAT0019983 | UAACAUAAUAGUGUGGAUUGA |
| hsa-miR-5694 | 480153_mir | MIMAT0022487 | CAGAUCAUGGGACUGUCUCAG |
| hsa-miR-5695 | 480154_mir | MIMAT0022488 | ACUCCAAGAAGAAUCUAGACAG |
| hsa-miR-5698 | 480157_mir | MIMAT0022491 | UGGGGGAGUGCAGUGAUUGUGG |
| hsa-miR-5700 | 480736_mir | MIMAT0022493 | UAAUGCAUUAAAUUAUUGAAGG |
| hsa-miR-6083 | 480180_mir | MIMAT0023708 | CUUAUAUCAGAGGCUGUGGG |
| hsa-miR-8060 | 480604_mir | MIMAT0030987 | CCAUGAAGCAGUGGGUAGGAGGAC |
| hsa-miR-8087 | 480628_mir | MIMAT0031014 | GAAGACUUCUUGGAUUACAGGGG |
| hsa-miR-10b-3p | 477868_mir | MIMAT0004556 | ACAGAUUCGAUUCUAGGGGAAU |
| hsa-miR-1237-5p | 479550_mir | MIMAT0022946 | CGGGGGCGGGGCCGAAGCGCG |
| hsa-miR-1266-5p | 479559_mir | MIMAT0005920 | CCUCAGGGCUGUAGAACAGGGCU |
| hsa-miR-1287-3p | 479566_mir | MIMAT0026738 | CUCUAGCCACAGAUGCAGUGAU |
| hsa-miR-1306-5p | 478701_mir | MIMAT0022726 | CCACCUCCCCUGCAAACGUCCA |
| hsa-miR-138-1-3p | 478311_mir | MIMAT0004607 | GCUACUUCACAACACCAGGGCC |
| hsa-miR-139-3p | 477906_mir | MIMAT0004552 | UGGAGACGCGGCCCUGUUGGAGU |
| hsa-miR-203a-3p | 478316_mir | MIMAT0000264 | GUGAAAUGUUUAGGACCACUAG |
| hsa-miR-20a-3p | 478317_mir | MIMAT0004493 | ACUGCAUUAUGAGCACUUAAAG |
| hsa-miR-2115-3p | 479588_mir | MIMAT0011159 | CAUCAGAAUUCAUGGAGGCUAG |

**Supplementary Table 3-5: TaqMan probe information and target sequences for qPCR-based miRNA analysis.**

| Marker | Probe ID | miRBase ID | Mature sequence |
|---|---|---|---|
| hsa-miR-3126-3p | 479614_mir | MIMAT0015377 | CAUCUGGCAUCCGUCACACAGA |
| hsa-miR-3130-5p | 479621_mir | MIMAT0014995 | UACCCAGUCUCCGGUGCAGCC |
| hsa-miR-3152-5p | 479638_mir | MIMAT0019207 | AUUGCCUCUGUUCUAACACAAG |
| hsa-miR-3609 | 479681_mir | MIMAT0017986 | CAAAGUGAUGAGUAAUACUGGCUG |
| hsa-miR-3617-3p | 479685_mir | MIMAT0022966 | CAUCAGCACCCUAUGUCCUUUCU |
| hsa-miR-365a-5p | 478842_mir | MIMAT0009199 | AGGGACUUUUGGGGGCAGAUGUG |
| hsa-miR-375-5p | 483110_mir | MIMAT0037313 | GCGACGAGCCCCUCGCACAAACC |
| hsa-miR-4661-5p | 479895_mir | MIMAT0019729 | AACUAGCUCUGUGGAUCCUGAC |
| hsa-miR-4777-3p | 480037_mir | MIMAT0019935 | AUACCUCAUCUAGAAUGCUGUA |
| hsa-miR-4778-5p | 480039_mir | MIMAT0019936 | AAUUCUGUAAAGGAAGAAGAGG |
| hsa-miR-4782-5p | 480043_mir | MIMAT0019944 | UUCUGGAUAUGAAGACAAUCAA |
| hsa-miR-4798-5p | 480062_mir | MIMAT0019974 | UUCGGUAUACUUUGUGAAUUGG |
| hsa-miR-514a-5p | 478974_mir | MIMAT0022702 | UACUCUGGAGAGUGACAAUCAUG |
| hsa-miR-5189-3p | 480103_mir | MIMAT0027088 | UGCCAACCGUCAGAGCCCAGA |
| hsa-miR-518a-3p | 478981_mir | MIMAT0002863 | GAAAGCGCUUCCCUUUGCUGGA |
| hsa-miR-550a-5p | 477852_mir | MIMAT0004800 | AGUGCCUGAGGGAGUAAGAGCCC |
| hsa-miR-552-5p | 479037_mir | MIMAT0026615 | GUUUAACCUUUUGCCUGUUGG |
| hsa-miR-5586-5p | 480131_mir | MIMAT0022287 | UAUCCAGCUUGUUACUAUAUGC |
| hsa-miR-5590-3p | 480138_mir | MIMAT0022300 | AAUAAAGUUCAUGUAUGGCAA |
| hsa-miR-615-5p | 478176_mir | MIMAT0004804 | GGGGGUCCCCGGUGCUCGGAUC |
| hsa-miR-6503-3p | 480196_mir | MIMAT0025463 | GGGACUAGGAUGCAGACCUCC |
| hsa-miR-653-3p | 479133_mir | MIMAT0026625 | UUCACUGGAGUUUGUUUCAAUA |
| hsa-miR-668-3p | 479151_mir | MIMAT0003881 | UGUCACUCGGCUCGGCCCACUAC |
| hsa-miR-6728-5p | 480237_mir | MIMAT0027357 | UUGGGAUGGUAGGACCAGAGGGG |
| hsa-miR-6738-5p | 480255_mir | MIMAT0027377 | CGAGGGGUAGAAGAGCACAGGGG |
| hsa-miR-6749-5p | 480272_mir | MIMAT0027398 | UCGGGCCUGGGGUUGGGGGAGC |
| hsa-miR-6755-3p | 480281_mir | MIMAT0027411 | UGUUGUCAUGUUUUUUCCCUAG |
| hsa-miR-6765-3p | 480299_mir | MIMAT0027431 | UCACCUGGCUGGCCCGCCCAG |
| hsa-miR-6781-3p | 480333_mir | MIMAT0027463 | UGCCUCUUUUCCACGGCCUCAG |
| hsa-miR-6785-5p | 480341_mir | MIMAT0027470 | UGGGAGGGCGUGGAUGAUGGUG |
| hsa-miR-6794-3p | 480356_mir | MIMAT0027489 | CUCACUCUCAGUCCCUCCCU |
| hsa-miR-6797-5p | 480361_mir | MIMAT0027494 | AGGAGGGAAGGGGCUGAGAACAGGA |
| hsa-miR-6811-5p | 480388_mir | MIMAT0027522 | AUGCAGGCCUGUGUACAGCACU |
| hsa-miR-6855-3p | 480465_mir | MIMAT0027611 | AGACUGACCUUCAACCCCACAG |
| hsa-miR-767-5p | 479176_mir | MIMAT0003882 | UGCACCAUGGUUGUCUGAGCAUG |
| hsa-miR-7847-3p | 480584_mir | MIMAT0030422 | CGUGGAGGACGAGGAGGAGGC |
| hsa-miR-885-3p | 479188_mir | MIMAT0004948 | AGGCAGCGGGGUGUAGUGGAUA |
| hsa-miR-889-5p | 479193_mir | MIMAT0026719 | AAUGGCUGUCCGUAGUAUGGUC |
| hsa-miR-96-3p | 479222_mir | MIMAT0004510 | AAUCAUGUGCAGUGCCAAUAUG |
| hsa-miR-486-5p | 478128_mir | MIMAT0002177 | UCCUGUACUGAGCUGCCCCGAG |
| cel-miR-39-3p | 478293_mir | - | - |

**Supplementary Table 4-1: Serum miRNA expression levels in NC, PC and BC samples (Batch 1).**

| miRNA | Non-cancer | Pancreatic cancer | Breast cancer | p-value (NC-PC) | p-value (NC-BC) |
|---|---|---|---|---|---|
| hsa-let-7a-5p | 2.73 ± 0.52 | 3.53 ± 0.60 | 4.27 ± 1.00 | 1.06E-05 | 3.24E-03 |
| hsa-let-7b-3p | 7.36 ± 0.59 | 7.85 ± 0.67 | 8.67 ± 0.68 | 9.44E-03 | 1.20E-04 |
| hsa-let-7b-5p | 6.36 ± 0.32 | 7.22 ± 0.63 | 8.07 ± 0.69 | 3.14E-07 | 4.36E-05 |
| hsa-let-7c-5p | 6.34 ± 0.50 | 7.35 ± 0.66 | 7.33 ± 0.82 | 3.73E-07 | 0.941 |
| hsa-let-7d-3p | 2.24 ± 0.48 | 2.59 ± 0.79 | 3.28 ± 0.81 | 0.0697 | 4.53E-03 |
| hsa-let-7e-5p | 7.33 ± 0.59 | 8.69 ± 0.91 | 8.86 ± 1.22 | 1.64E-07 | 0.593 |
| hsa-let-7f-5p | 3.91 ± 0.67 | 4.94 ± 0.66 | 5.57 ± 1.19 | 2.27E-06 | 0.0293 |
| hsa-let-7g-5p | 3.88 ± 0.46 | 5.01 ± 0.54 | 5.85 ± 1.04 | 6.52E-10 | 9.86E-04 |
| hsa-let-7i-5p | 1.09 ± 0.31 | 2.05 ± 0.39 | 2.85 ± 0.79 | 2.41E-12 | 5.38E-05 |
| hsa-miR-101-3p | -0.40 ± 0.54 | 1.13 ± 0.51 | 1.34 ± 0.61 | 3.54E-13 | 0.187 |
| hsa-miR-103a-3p | 1.19 ± 0.48 | 2.60 ± 0.67 | 3.04 ± 1.03 | 9.60E-11 | 0.0813 |
| hsa-miR-106b-5p | 3.63 ± 0.41 | 5.07 ± 0.65 | 6.58 ± 1.46 | 5.47E-12 | 2.92E-05 |
| hsa-miR-107 | 1.27 ± 0.41 | 2.37 ± 0.41 | 2.99 ± 0.73 | 4.05E-12 | 7.64E-04 |
| hsa-miR-10b-5p | 6.50 ± 0.77 | 7.03 ± 0.79 | 7.35 ± 0.78 | 0.0221 | 0.163 |
| hsa-miR-122-5p | 2.64 ± 2.00 | 2.67 ± 1.69 | 5.64 ± 1.49 | 0.953 | 5.84E-08 |
| hsa-miR-1229-3p | 7.98 ± 1.04 | 9.25 ± 1.74 | 8.64 ± 1.53 | 4.06E-03 | 0.243 |
| hsa-miR-1246 | 2.59 ± 1.30 | 4.09 ± 1.43 | 4.79 ± 1.71 | 4.20E-04 | 0.133 |
| hsa-miR-1249-3p | 6.65 ± 0.59 | 7.53 ± 1.40 | 7.36 ± 1.51 | 6.78E-03 | 0.685 |
| hsa-miR-125a-5p | 3.73 ± 0.55 | 4.56 ± 0.97 | 4.59 ± 1.24 | 6.78E-04 | 0.914 |
| hsa-miR-125b-5p | 5.62 ± 2.53 | 4.30 ± 1.72 | 8.42 ± 3.53 | 0.0399 | 5.59E-06 |
| hsa-miR-126-3p | 1.13 ± 0.56 | 2.25 ± 0.62 | 2.89 ± 1.04 | 4.70E-08 | 0.0130 |
| hsa-miR-127-3p | 7.43 ± 0.67 | 9.57 ± 1.23 | 9.21 ± 1.72 | 1.77E-09 | 0.408 |
| hsa-miR-128-3p | 3.85 ± 0.56 | 4.62 ± 0.79 | 4.59 ± 1.12 | 3.62E-04 | 0.934 |
| hsa-miR-1296-5p | 7.63 ± 0.62 | 8.91 ± 1.13 | 8.78 ± 1.39 | 1.52E-05 | 0.724 |
| hsa-miR-1307-3p | 7.85 ± 0.45 | 8.51 ± 0.85 | 9.16 ± 1.09 | 1.59E-03 | 0.0246 |
| hsa-miR-132-3p | 3.55 ± 0.84 | 5.31 ± 1.21 | 5.43 ± 1.19 | 5.11E-07 | 0.716 |
| hsa-miR-133a-3p | 8.68 ± 1.01 | 8.61 ± 1.53 | 10.01 ± 1.12 | 0.863 | 6.92E-04 |
| hsa-miR-1-3p | 3.45 ± 0.79 | 4.54 ± 1.07 | 5.08 ± 0.98 | 2.21E-04 | 0.0740 |
| hsa-miR-140-3p | 3.65 ± 0.60 | 4.00 ± 0.69 | 3.92 ± 0.78 | 0.0667 | 0.698 |
| hsa-miR-142-3p | 0.42 ± 0.54 | 3.04 ± 0.64 | 3.12 ± 1.17 | 1.24E-19 | 0.754 |
| hsa-miR-142-5p | -1.80 ± 0.57 | 0.41 ± 0.80 | 1.65 ± 1.36 | 1.64E-14 | 3.70E-04 |
| hsa-miR-143-3p | 2.26 ± 0.81 | 3.56 ± 1.08 | 4.67 ± 1.15 | 2.29E-05 | 1.25E-03 |
| hsa-miR-146a-5p | 1.38 ± 0.61 | 2.28 ± 0.61 | 3.03 ± 0.95 | 7.50E-06 | 2.16E-03 |
| hsa-miR-148a-3p | 1.15 ± 0.76 | 1.99 ± 0.87 | 3.18 ± 0.87 | 8.37E-04 | 2.17E-05 |
| hsa-miR-148b-3p | 3.69 ± 0.59 | 4.67 ± 0.83 | 4.57 ± 1.13 | 2.38E-05 | 0.737 |
| hsa-miR-150-5p | 2.53 ± 0.63 | 3.71 ± 0.67 | 3.57 ± 0.63 | 1.05E-07 | 0.461 |
| hsa-miR-151a-3p | 3.15 ± 0.59 | 4.11 ± 0.92 | 4.15 ± 1.19 | 9.63E-05 | 0.883 |
| hsa-miR-155-5p | 4.89 ± 0.80 | 6.22 ± 0.72 | 6.65 ± 1.26 | 2.27E-07 | 0.151 |
| hsa-miR-15a-5p | -0.17 ± 0.53 | 0.39 ± 0.73 | 0.36 ± 0.61 | 3.84E-03 | 0.874 |
| hsa-miR-15b-5p | -1.51 ± 0.58 | -0.29 ± 1.06 | -0.27 ± 1.13 | 1.11E-05 | 0.968 |
| hsa-miR-16-2-3p | 4.81 ± 0.27 | 5.66 ± 0.31 | 6.58 ± 0.64 | 2.30E-13 | 8.41E-08 |
| hsa-miR-16-5p | -1.10 ± 0.29 | -0.54 ± 0.34 | 0.44 ± 0.71 | 2.06E-07 | 2.06E-07 |
| hsa-miR-17-5p | 0.85 ± 0.30 | 1.62 ± 0.43 | 2.37 ± 0.79 | 5.17E-09 | 1.66E-04 |
| hsa-miR-181a-5p | 3.15 ± 0.67 | 4.27 ± 0.62 | 4.81 ± 1.16 | 2.83E-07 | 0.0498 |
| hsa-miR-181b-5p | 7.49 ± 0.52 | 8.51 ± 0.80 | 8.96 ± 0.92 | 4.16E-06 | 0.0736 |
| hsa-miR-186-5p | 2.55 ± 0.46 | 3.62 ± 0.53 | 4.48 ± 0.87 | 2.19E-09 | 1.42E-04 |
| hsa-miR-18a-5p | 3.79 ± 0.44 | 4.91 ± 0.50 | 5.53 ± 0.98 | 1.27E-10 | 8.24E-03 |
| hsa-miR-190a-5p | 7.49 ± 0.35 | 9.59 ± 0.66 | 10.16 ± 1.08 | 6.17E-18 | 0.0304 |
| hsa-miR-191-5p | 1.32 ± 0.52 | 2.33 ± 0.71 | 2.94 ± 1.02 | 9.47E-07 | 0.0211 |
| hsa-miR-192-5p | 4.41 ± 0.59 | 4.38 ± 0.99 | 4.91 ± 1.15 | 0.912 | 0.0924 |
| hsa-miR-193a-5p | 6.30 ± 1.28 | 7.36 ± 0.85 | 8.95 ± 0.88 | 1.54E-03 | 8.29E-08 |

**Supplementary Table 4-2: Serum miRNA expression levels in NC, PC and BC samples (Batch 1).**

| miRNA | Non-cancer | Pancreatic cancer | Breast cancer | p-value (NC-PC) | p-value (NC-BC) |
|---|---|---|---|---|---|
| hsa-miR-193b-5p | 12.57 ± 3.45 | 10.29 ± 3.39 | 11.93 ± 3.56 | 0.0579 | 0.181 |
| hsa-miR-194-5p | 5.80 ± 0.85 | 6.94 ± 0.86 | 8.67 ± 0.72 | 3.23E-05 | 1.49E-09 |
| hsa-miR-196a-5p | 7.89 ± 1.01 | 9.81 ± 1.29 | 10.11 ± 1.11 | 6.55E-07 | 0.401 |
| hsa-miR-196b-5p | 10.97 ± 0.92 | 11.48 ± 0.66 | 11.96 ± 1.12 | 0.0328 | 0.0768 |
| hsa-miR-197-3p | 3.10 ± 0.60 | 3.79 ± 0.94 | 4.54 ± 0.88 | 4.39E-03 | 6.30E-03 |
| hsa-miR-199a-3p | 1.31 ± 0.59 | 2.09 ± 0.77 | 2.80 ± 1.12 | 2.98E-04 | 0.0132 |
| hsa-miR-19b-3p | -0.44 ± 0.40 | 1.07 ± 0.42 | 1.92 ± 0.78 | 1.27E-16 | 2.48E-05 |
| hsa-miR-2052 | 13.67 ± 1.33 | 15.08 ± 1.70 | 14.52 ± 2.04 | 0.0103 | 0.416 |
| hsa-miR-205-5p | 2.92 ± 0.64 | 4.83 ± 1.07 | 4.76 ± 0.83 | 1.60E-09 | 0.814 |
| hsa-miR-206 | 10.45 ± 2.12 | 9.71 ± 2.10 | 9.89 ± 2.23 | 0.239 | 0.783 |
| hsa-miR-20a-5p | 1.44 ± 0.32 | 2.17 ± 0.40 | 2.86 ± 0.77 | 7.89E-09 | 3.59E-04 |
| hsa-miR-21-5p | -21.04 ± 0.50 | -0.42 ± 0.65 | 0.15 ± 1.00 | 1.12E-59 | 0.0245 |
| hsa-miR-215-5p | 6.30 ± 0.90 | 6.25 ± 1.10 | 6.85 ± 1.54 | 0.870 | 0.126 |
| hsa-miR-221-3p | -0.01 ± 0.49 | 1.01 ± 0.74 | 1.61 ± 0.86 | 9.26E-07 | 0.0126 |
| hsa-miR-222-3p | 4.28 ± 0.52 | 5.40 ± 0.60 | 6.00 ± 0.85 | 1.05E-08 | 7.51E-03 |
| hsa-miR-223-3p | -5.42 ± 0.70 | -4.55 ± 1.05 | -3.49 ± 1.00 | 1.54E-03 | 8.17E-04 |
| hsa-miR-223-5p | 5.92 ± 1.27 | 6.74 ± 1.52 | 8.37 ± 1.49 | 0.0468 | 5.16E-04 |
| hsa-miR-22-3p | -3.66 ± 0.71 | -2.02 ± 0.63 | -1.38 ± 0.89 | 5.61E-11 | 5.95E-03 |
| hsa-miR-22-5p | 4.55 ± 0.66 | 5.89 ± 0.63 | 6.87 ± 0.90 | 5.39E-09 | 6.93E-05 |
| hsa-miR-2276-3p | 10.84 ± 1.34 | 11.36 ± 1.24 | 12.61 ± 1.61 | 0.178 | 0.0161 |
| hsa-miR-23a-3p | 2.46 ± 0.60 | 3.23 ± 0.77 | 3.94 ± 0.92 | 3.54E-04 | 5.50E-03 |
| hsa-miR-23b-3p | 1.34 ± 0.57 | 2.07 ± 0.71 | 2.71 ± 0.91 | 2.82E-04 | 8.68E-03 |
| hsa-miR-24-3p | 0.82 ± 0.47 | 1.68 ± 0.79 | 2.76 ± 0.98 | 4.19E-05 | 1.08E-04 |
| hsa-miR-25-3p | -0.10 ± 0.60 | 0.29 ± 0.40 | 1.22 ± 0.42 | 0.0122 | 5.77E-10 |
| hsa-miR-26b-5p | 0.71 ± 0.51 | 2.02 ± 0.57 | 2.73 ± 1.11 | 8.86E-11 | 7.82E-03 |
| hsa-miR-27a-3p | 2.09 ± 0.63 | 2.88 ± 0.72 | 3.69 ± 1.07 | 2.20E-04 | 3.28E-03 |
| hsa-miR-27b-3p | 2.04 ± 0.58 | 3.04 ± 0.69 | 3.65 ± 0.95 | 2.06E-06 | 0.0142 |
| hsa-miR-27b-5p | 10.08 ± 1.97 | 11.78 ± 1.70 | 11.97 ± 2.26 | 2.39E-03 | 0.746 |
| hsa-miR-28-3p | 7.04 ± 0.60 | 7.96 ± 0.73 | 8.46 ± 0.91 | 2.04E-05 | 0.0429 |
| hsa-miR-29a-3p | 0.67 ± 0.73 | 2.49 ± 0.83 | 3.50 ± 0.79 | 1.99E-10 | 8.35E-05 |
| hsa-miR-29c-3p | 0.00 ± 0.60 | 1.78 ± 0.54 | 2.57 ± 0.79 | 2.91E-14 | 1.95E-04 |
| hsa-miR-30d-5p | 1.88 ± 0.52 | 2.59 ± 0.58 | 3.19 ± 0.63 | 4.40E-05 | 1.27E-03 |
| hsa-miR-30e-5p | 4.72 ± 0.69 | 6.84 ± 1.52 | 8.96 ± 3.24 | 1.35E-07 | 5.73E-03 |
| hsa-miR-3151-5p | 7.09 ± 1.03 | 8.31 ± 1.23 | 8.64 ± 1.83 | 6.12E-04 | 0.469 |
| hsa-miR-3155a | 5.45 ± 1.28 | 6.70 ± 1.47 | 6.67 ± 2.05 | 2.98E-03 | 0.945 |
| hsa-miR-31-5p | 9.49 ± 1.10 | 11.93 ± 0.88 | 11.80 ± 1.11 | 5.97E-11 | 0.643 |
| hsa-miR-3168 | 8.08 ± 0.95 | 8.93 ± 1.16 | 8.88 ± 1.74 | 7.71E-03 | 0.900 |
| hsa-miR-3178 | 6.92 ± 0.68 | 6.55 ± 0.82 | 7.22 ± 0.74 | 0.100 | 5.07E-03 |
| hsa-miR-3182 | 3.70 ± 1.17 | 4.48 ± 1.51 | 4.67 ± 1.89 | 0.0498 | 0.713 |
| hsa-miR-320a-3p | 2.34 ± 0.54 | 3.02 ± 0.47 | 3.59 ± 0.55 | 2.91E-05 | 4.24E-04 |
| hsa-miR-320b | 4.30 ± 0.67 | 5.11 ± 0.47 | 5.71 ± 0.57 | 1.39E-05 | 2.39E-04 |
| hsa-miR-324-3p | 6.29 ± 0.38 | 7.34 ± 0.67 | 7.41 ± 0.77 | 2.81E-08 | 0.750 |
| hsa-miR-324-5p | 5.01 ± 0.41 | 6.15 ± 0.59 | 6.76 ± 0.96 | 5.80E-10 | 0.0117 |
| hsa-miR-32-5p | 1.92 ± 0.81 | 3.19 ± 1.31 | 2.68 ± 1.45 | 2.07E-04 | 0.209 |
| hsa-miR-326 | 4.64 ± 0.59 | 5.25 ± 0.70 | 5.21 ± 1.01 | 2.34E-03 | 0.872 |
| hsa-miR-328-3p | 6.24 ± 0.53 | 7.00 ± 0.81 | 7.25 ± 0.95 | 3.61E-04 | 0.317 |
| hsa-miR-329-5p | 6.32 ± 1.21 | 7.67 ± 1.60 | 7.67 ± 2.17 | 1.89E-03 | 0.988 |
| hsa-miR-338-5p | 8.02 ± 0.98 | 9.19 ± 1.56 | 9.21 ± 1.67 | 3.23E-03 | 0.969 |
| hsa-miR-339-5p | 3.82 ± 0.45 | 5.43 ± 0.62 | 5.90 ± 1.15 | 1.33E-13 | 0.0841 |
| hsa-miR-342-3p | 4.40 ± 0.52 | 5.13 ± 0.61 | 5.30 ± 0.63 | 5.83E-05 | 0.328 |
| hsa-miR-34a-5p | 6.39 ± 1.52 | 7.43 ± 1.39 | 10.32 ± 1.67 | 0.0169 | 6.42E-08 |

**Supplementary Table 4-3: Serum miRNA expression levels in NC, PC and BC samples (Batch 1).**

| miRNA | Non-cancer | Pancreatic cancer | Breast cancer | p-value (NC-PC) | p-value (NC-BC) |
|---|---|---|---|---|---|
| hsa-miR-3613-3p | 6.83 ± 0.54 | 7.48 ± 0.91 | 8.49 ± 0.92 | 4.79E-03 | 3.86E-04 |
| hsa-miR-3613-5p | 12.22 ± 2.42 | 9.05 ± 1.84 | 10.85 ± 1.75 | 0.0118 | 0.0573 |
| hsa-miR-3615 | 3.69 ± 0.67 | 4.17 ± 0.95 | 4.44 ± 1.06 | 0.0505 | 0.356 |
| hsa-miR-361-5p | 3.79 ± 0.54 | 4.18 ± 0.87 | 5.27 ± 0.87 | 0.0702 | 8.03E-05 |
| hsa-miR-375-3p | 5.08 ± 0.93 | 8.37 ± 2.03 | 9.44 ± 3.24 | 4.27E-09 | 0.175 |
| hsa-miR-378a-3p | 6.45 ± 0.81 | 7.49 ± 0.66 | 8.76 ± 0.80 | 1.41E-05 | 2.58E-07 |
| hsa-miR-382-5p | 7.22 ± 0.81 | 8.93 ± 1.19 | 8.69 ± 1.68 | 4.78E-07 | 0.560 |
| hsa-miR-409-3p | 5.38 ± 1.11 | 7.17 ± 1.75 | 6.89 ± 1.55 | 1.11E-04 | 0.561 |
| hsa-miR-423-3p | 4.27 ± 0.50 | 5.18 ± 0.81 | 5.62 ± 0.95 | 2.54E-05 | 0.0908 |
| hsa-miR-423-5p | 2.55 ± 0.40 | 2.95 ± 0.66 | 3.85 ± 0.57 | 0.0153 | 6.97E-06 |
| hsa-miR-424-3p | 6.75 ± 0.81 | 7.29 ± 1.01 | 7.32 ± 1.04 | 0.0454 | 0.929 |
| hsa-miR-425-3p | 6.08 ± 0.47 | 6.82 ± 0.78 | 7.49 ± 0.95 | 2.58E-04 | 0.0102 |
| hsa-miR-425-5p | 1.85 ± 0.55 | 2.61 ± 0.54 | 3.40 ± 0.60 | 2.07E-05 | 1.85E-05 |
| hsa-miR-4300 | 5.43 ± 0.81 | 6.36 ± 1.06 | 6.13 ± 1.52 | 1.40E-03 | 0.549 |
| hsa-miR-4302 | 14.50 ± 0.99 | 15.74 ± 1.07 | 16.25 ± 0.83 | 1.32E-04 | 0.0745 |
| hsa-miR-432-5p | 11.10 ± 2.49 | 11.94 ± 2.47 | 10.64 ± 2.98 | 0.270 | 0.124 |
| hsa-miR-433-3p | 5.30 ± 1.03 | 7.03 ± 1.91 | 7.25 ± 1.81 | 3.18E-04 | 0.687 |
| hsa-miR-4433a-3p | 8.87 ± 1.10 | 9.84 ± 0.90 | 11.01 ± 1.94 | 1.96E-03 | 0.0137 |
| hsa-miR-4433b-5p | 6.97 ± 3.02 | 6.75 ± 1.38 | 6.49 ± 2.08 | 0.750 | 0.624 |
| hsa-miR-4446-3p | 7.45 ± 0.71 | 8.57 ± 1.33 | 8.73 ± 1.48 | 6.52E-04 | 0.702 |
| hsa-miR-4466 | 7.45 ± 0.71 | 8.57 ± 1.33 | 8.73 ± 1.48 | 6.52E-04 | 0.702 |
| hsa-miR-451a | -5.65 ± 0.41 | -3.75 ± 0.41 | -3.16 ± 0.56 | 2.08E-20 | 1.41E-04 |
| hsa-miR-4732-5p | 12.06 ± 0.86 | 11.62 ± 0.94 | 12.07 ± 0.67 | 0.0949 | 0.0587 |
| hsa-miR-4770 | 3.00 ± 0.96 | 3.73 ± 1.35 | 3.86 ± 1.44 | 0.0360 | 0.746 |
| hsa-miR-483-3p | 8.04 ± 1.32 | 8.70 ± 0.86 | 9.65 ± 0.58 | 0.0474 | 5.05E-05 |
| hsa-miR-483-5p | 9.05 ± 2.03 | 9.78 ± 1.45 | 11.91 ± 1.42 | 0.157 | 5.41E-06 |
| hsa-miR-484 | 1.40 ± 0.32 | 1.64 ± 0.55 | 1.96 ± 0.60 | 0.0727 | 0.0590 |
| hsa-miR-485-3p | 6.34 ± 1.15 | 8.51 ± 1.85 | 7.93 ± 1.60 | 1.34E-05 | 0.250 |
| hsa-miR-485-5p | 9.52 ± 1.34 | 9.47 ± 1.60 | 9.89 ± 1.65 | 0.901 | 0.370 |
| hsa-miR-486-3p | 6.78 ± 0.57 | 7.02 ± 0.54 | 7.60 ± 0.54 | 0.139 | 5.82E-04 |
| hsa-miR-493-3p | 9.45 ± 1.31 | 11.26 ± 2.53 | 11.27 ± 2.07 | 3.54E-03 | 0.986 |
| hsa-miR-5007-3p | 10.92 ± 0.78 | 12.63 ± 0.99 | 12.69 ± 0.99 | 3.16E-08 | 0.836 |
| hsa-miR-505-5p | 9.24 ± 0.92 | 9.35 ± 0.72 | 10.71 ± 1.47 | 0.648 | 1.77E-04 |
| hsa-miR-513-5p | 16.14 ± 1.12 | 17.33 ± 1.31 | 17.46 ± 1.84 | 2.24E-03 | 0.782 |
| hsa-miR-520e-3p | 8.36 ± 1.24 | 9.15 ± 1.27 | 8.84 ± 1.15 | 0.0334 | 0.379 |
| hsa-miR-532-5p | 6.14 ± 0.32 | 6.97 ± 0.39 | 8.10 ± 0.66 | 2.73E-10 | 4.50E-09 |
| hsa-miR-542-3p | 6.47 ± 1.17 | 7.35 ± 1.46 | 7.03 ± 1.56 | 0.0251 | 0.471 |
| hsa-miR-559 | 7.51 ± 1.15 | 8.48 ± 1.42 | 8.24 ± 1.70 | 0.0127 | 0.610 |
| hsa-miR-574-3p | 14.00 ± 2.22 | 11.74 ± 2.02 | 13.22 ± 1.96 | 6.85E-04 | 0.0142 |
| hsa-miR-574-5p | 4.98 ± 0.73 | 6.80 ± 1.07 | 7.17 ± 1.23 | 1.45E-08 | 0.267 |
| hsa-miR-576-3p | 5.66 ± 0.79 | 6.10 ± 1.12 | 6.26 ± 1.25 | 0.119 | 0.648 |
| hsa-miR-584-5p | 0.75 ± 0.49 | 1.77 ± 0.81 | 2.01 ± 0.83 | 3.66E-06 | 0.311 |
| hsa-miR-606 | 15.57 ± 1.05 | 16.72 ± 0.90 | 17.23 ± 0.91 | 1.72E-04 | 0.0592 |
| hsa-miR-616-3p | 11.84 ± 1.51 | 12.90 ± 2.38 | 12.83 ± 2.30 | 0.101 | 0.927 |
| hsa-miR-625-3p | 9.28 ± 1.15 | 9.23 ± 0.92 | 9.52 ± 1.24 | 0.865 | 0.354 |
| hsa-miR-629-5p | 7.17 ± 0.50 | 7.79 ± 0.86 | 8.91 ± 0.77 | 3.90E-03 | 1.97E-05 |
| hsa-miR-652-3p | 2.37 ± 0.62 | 3.33 ± 1.05 | 3.52 ± 1.03 | 3.28E-04 | 0.539 |
| hsa-miR-660-5p | 4.90 ± 0.49 | 6.19 ± 0.52 | 7.24 ± 0.78 | 1.66E-11 | 1.63E-06 |
| hsa-miR-664a-3p | 7.09 ± 0.68 | 7.60 ± 0.70 | 7.89 ± 1.03 | 0.0147 | 0.259 |
| hsa-miR-665 | 8.62 ± 1.86 | 10.20 ± 2.37 | 10.97 ± 3.29 | 0.0159 | 0.378 |
| hsa-miR-6756-5p | 3.84 ± 0.74 | 4.72 ± 1.05 | 5.14 ± 1.34 | 1.60E-03 | 0.235 |

**Supplementary Table 4-4: Serum miRNA expression levels in NC, PC and BC samples (Batch 1).**

| miRNA | Non-cancer | Pancreatic cancer | Breast cancer | p-value (NC-PC) | p-value (NC-BC) |
|---|---|---|---|---|---|
| hsa-miR-6859-5p | 12.58 ± 1.22 | 13.55 ± 1.38 | 13.25 ± 1.62 | 0.0130 | 0.487 |
| hsa-miR-6872-5p | 11.96 ± 1.19 | 12.63 ± 1.54 | 13.36 ± 1.85 | 0.109 | 0.157 |
| hsa-miR-6886-3p | 10.55 ± 0.99 | 12.07 ± 1.43 | 11.92 ± 2.18 | 9.53E-05 | 0.777 |
| hsa-miR-744-5p | 5.03 ± 0.59 | 6.02 ± 0.70 | 6.63 ± 1.16 | 3.70E-06 | 0.0312 |
| hsa-miR-758-3p | 5.82 ± 1.09 | 8.08 ± 2.23 | 7.57 ± 1.77 | 5.47E-05 | 0.392 |
| hsa-miR-769-5p | 7.64 ± 0.94 | 8.58 ± 1.16 | 8.64 ± 1.44 | 3.33E-03 | 0.868 |
| hsa-miR-885-5p | 7.44 ± 1.97 | 7.16 ± 1.51 | 9.67 ± 1.21 | 0.585 | 8.78E-08 |
| hsa-miR-92b-3p | 1.20 ± 0.30 | 1.34 ± 0.38 | 1.78 ± 0.42 | 0.159 | 3.97E-04 |
| hsa-miR-93-3p | 5.87 ± 0.45 | 6.68 ± 0.78 | 6.61 ± 0.93 | 6.06E-05 | 0.776 |
| hsa-miR-93-5p | 1.26 ± 0.55 | 1.94 ± 0.53 | 2.80 ± 0.66 | 8.37E-05 | 1.06E-05 |
| hsa-miR-941 | 5.74 ± 0.91 | 6.91 ± 1.29 | 8.29 ± 0.78 | 6.62E-04 | 5.12E-05 |
| hsa-miR-9-5p | 6.42 ± 0.73 | 7.60 ± 1.05 | 7.70 ± 1.60 | 4.38E-05 | 0.795 |
| hsa-miR-98-5p | 4.14 ± 0.80 | 4.84 ± 0.89 | 5.67 ± 1.39 | 5.87E-03 | 0.0175 |
| hsa-miR-99a-5p | 6.08 ± 0.69 | 6.02 ± 0.99 | 6.61 ± 1.51 | 0.820 | 0.116 |
| hsa-miR-99b-5p | 6.24 ± 0.57 | 7.42 ± 0.88 | 7.35 ± 1.03 | 1.32E-06 | 0.779 |

**Supplementary Table 5: Serum miRNA expression levels in NC, PC and other cancer samples (Batch 2).**

| miRNA | NC | | PC | | Other cancers | | F | p-value |
|---|---|---|---|---|---|---|---|---|
| | AVG | SD | AVG | SD | AVG | SD | | |
| hsa-miR-106b-5p | 0.222 | 0.0740 | 0.0883 | 0.0486 | 0.117 | 0.0752 | 87.5 | 1.01E-33 |
| hsa-miR-122-5p | 0.611 | 0.793 | 0.289 | 0.387 | 0.195 | 0.753 | 11.3 | 1.52E-05 |
| hsa-miR-125a-5p | 0.0587 | 0.0253 | 0.0286 | 0.0213 | 0.0361 | 0.0317 | 24.8 | 4.75E-11 |
| hsa-miR-125b-5p | 0.0463 | 0.0312 | 0.0266 | 0.0187 | 0.0256 | 0.0465 | 8.49 | 2.35E-04 |
| hsa-miR-128-3p | 0.0272 | 0.0115 | 0.0131 | 0.00673 | 0.0168 | 0.0107 | 44.2 | 1.61E-18 |
| hsa-miR-142-3p | 2.65 | 1.13 | 0.706 | 0.428 | 0.993 | 0.702 | 185 | 3.33E-62 |
| hsa-miR-143-3p | 0.0861 | 0.0489 | 0.0285 | 0.0225 | 0.0463 | 0.0497 | 34.2 | 9.75E-15 |
| hsa-miR-151a-3p | 0.0299 | 0.0147 | 0.0130 | 0.0101 | 0.0157 | 0.0109 | 58.1 | 1.31E-23 |
| hsa-miR-190a-5p | 0.0168 | 0.00503 | 0.00512 | 0.00434 | 0.00616 | 0.00401 | 228 | 1.05E-72 |
| hsa-miR-193a-5p | 0.0408 | 0.0332 | 0.0125 | 0.0108 | 0.0133 | 0.0164 | 72.0 | 1.69E-28 |
| hsa-miR-194-5p | 0.0569 | 0.0359 | 0.0243 | 0.0159 | 0.0231 | 0.0626 | 13.0 | 2.92E-06 |
| hsa-miR-205-5p | 0.0198 | 0.00861 | 0.00735 | 0.00664 | 0.0141 | 0.0649 | 1.06 | 0.348 |
| hsa-miR-223-5p | 0.0168 | 0.00923 | 0.00468 | 0.00416 | 0.00737 | 0.00804 | 59.4 | 4.34E-24 |
| hsa-miR-29a-3p | 0.361 | 0.172 | 0.127 | 0.0903 | 0.150 | 0.152 | 74.0 | 3.64E-29 |
| hsa-miR-29c-3p | 0.551 | 0.205 | 0.258 | 0.114 | 0.313 | 0.199 | 61.5 | 7.75E-25 |
| hsa-miR-30e-5p | 0.754 | 0.276 | 0.279 | 0.164 | 0.331 | 0.227 | 127 | 4.25E-46 |
| hsa-miR-324-3p | 0.00759 | 0.00341 | 0.00346 | 0.00182 | 0.00451 | 0.00297 | 48.3 | 4.93E-20 |
| hsa-miR-34a-5p | 0.0192 | 0.0211 | 0.0118 | 0.0115 | 0.00668 | 0.0127 | 28.1 | 2.34E-12 |
| hsa-miR-3613-3p | 0.00640 | 0.00246 | 0.00254 | 0.00139 | 0.00358 | 0.00302 | 45.7 | 4.36E-19 |
| hsa-miR-3613-5p | 0.0735 | 0.0237 | 0.0274 | 0.0124 | 0.0345 | 0.0241 | 111 | 2.34E-41 |
| hsa-miR-375-3p | 0.0162 | 0.00835 | 0.00532 | 0.0178 | 0.00613 | 0.0281 | 5.96 | 2.75E-03 |
| hsa-miR-4433a-3p | 0.00446 | 0.00305 | 0.00128 | 0.00109 | 0.00176 | 0.00168 | 77.9 | 1.71E-30 |
| hsa-miR-4770 | 0.0195 | 0.0122 | 0.00591 | 0.00508 | 0.00959 | 0.0102 | 43.1 | 4.00E-18 |
| hsa-miR-483-5p | 0.00790 | 0.00839 | 0.00448 | 0.00427 | 0.00353 | 0.0102 | 7.37 | 6.93E-04 |
| hsa-miR-574-3p | 0.0198 | 0.00782 | 0.00882 | 0.00639 | 0.0101 | 0.0101 | 39.6 | 8.65E-17 |
| hsa-miR-885-5p | 0.0150 | 0.0185 | 0.00659 | 0.00794 | 0.00388 | 0.0138 | 21.4 | 1.12E-09 |
| hsa-miR-941 | 0.00352 | 0.00151 | 0.00109 | 0.00106 | 0.00178 | 0.00213 | 36.6 | 1.19E-15 |

(NC, non-cancer; PC, pancreatic cancer; AVG, average; SD, standard deviation)

**Supplementary Table 6: Serum miRNA expression levels in NC, PC and other cancer samples (Batch 3).**

| miRNA | NC | | PC | | Other cancers | | F | p-value |
|---|---|---|---|---|---|---|---|---|
| | AVG | SD | AVG | SD | AVG | SD | | |
| hsa-miR-142-3p | 1.81 | 0.937 | 0.569 | 0.547 | 0.744 | 0.695 | 62.6 | 2.80E-24 |
| hsa-miR-190a-5p | 0.0155 | 0.00815 | 0.00540 | 0.00536 | 0.00573 | 0.00430 | 91 | 3.90E-33 |
| hsa-miR-194-5p | 0.0490 | 0.0298 | 0.0411 | 0.0460 | 0.0231 | 0.0178 | 30.3 | 5.90E-13 |
| hsa-miR-205-5p | 0.0260 | 0.0172 | 0.00954 | 0.00923 | 0.0281 | 0.215 | 0.3 | 0.746 |
| hsa-miR-223-5p | 0.0238 | 0.0171 | 0.00645 | 0.00699 | 0.00980 | 0.0123 | 37.4 | 1.40E-15 |
| hsa-miR-29c-3p | 0.639 | 0.323 | 0.323 | 0.184 | 0.404 | 0.288 | 23.3 | 2.80E-10 |
| hsa-miR-30e-5p | 1.02 | 0.527 | 0.411 | 0.222 | 0.522 | 0.379 | 50.3 | 3.70E-20 |
| hsa-miR-324-3p | 0.0109 | 0.00456 | 0.00433 | 0.00266 | 0.00564 | 0.00447 | 46.2 | 1.00E-18 |
| hsa-miR-34a-5p | 0.0143 | 0.0118 | 0.0172 | 0.0271 | 0.00789 | 0.00791 | 15.1 | 4.80E-07 |
| hsa-miR-3613-3p | 0.0719 | 0.513 | 0.00266 | 0.00188 | 0.00365 | 0.00362 | 3 | 0.0533 |
| hsa-miR-3613-5p | 0.123 | 0.0635 | 0.0484 | 0.0242 | 0.0596 | 0.0366 | 70.1 | 1.00E-26 |
| hsa-miR-483-5p | 0.00446 | 0.00460 | 0.00638 | 0.00903 | 0.00280 | 0.00596 | 8.8 | 1.80E-04 |
| hsa-miR-885-5p | 0.00590 | 0.0103 | 0.0138 | 0.0221 | 0.00310 | 0.00566 | 25.1 | 5.80E-11 |

(NC, non-cancer; PC, pancreatic cancer; AVG, average; SD, standard deviation)

**Supplementary Table 7: Serum miRNA expression levels in NC, PC and other cancer samples (Batch 4).**

| miRNA | NC | | PC | | Other cancers | | F | p-value |
|---|---|---|---|---|---|---|---|---|
| | AVG | SD | AVG | SD | AVG | SD | | |
| hsa-miR-205-5p | 0.514 | 0.174 | 0.298 | 0.229 | 0.471 | 0.905 | 6.4 | 1.70E-03 |
| hsa-miR-29c-3p | 14.216 | 4.423 | 12.532 | 4.384 | 16.224 | 5.468 | 59.2 | 1.70E-25 |
| hsa-miR-324-3p | 0.238 | 0.103 | 0.168 | 0.0632 | 0.230 | 0.0888 | 57.6 | 7.60E-25 |
| hsa-miR-34a-5p | 0.362 | 0.251 | 0.522 | 0.325 | 0.286 | 0.180 | 119.5 | 5.30E-49 |
| hsa-miR-483-5p | 0.134 | 0.101 | 0.185 | 0.180 | 0.101 | 0.112 | 49.3 | 1.70E-21 |
| hsa-miR-486-5p | 23.895 | 12.671 | 42.926 | 27.385 | 52.228 | 32.963 | 88.1 | 5.60E-37 |

(NC, non-cancer; PC, pancreatic cancer; AVG, average; SD, standard deviation)
